# Temporal loss of genome-wide and immunogenetic diversity in a near-extinct parrot

**DOI:** 10.1101/2024.11.10.622863

**Authors:** Luke W. Silver, Katherine A. Farquharson, Emma Peel, M. Thomas P. Gilbert, Katherine Belov, Hernán E. Morales, Carolyn J. Hogg

## Abstract

Loss of genetic diversity threatens a species’ adaptive potential and long-term resilience. Predicted to be extinct by 2038, the orange-bellied parrot (*Neophema chrysogaster*) is a Critically Endangered migratory bird threatened by numerous viral, bacterial and fungal diseases. The species has undergone multiple population crashes, reaching a low of three wild-born females and 13 males in 2016 and is now represented by only a single wild population and individuals in the captive breeding program. Here we used our high-quality long-read reference genome, and contemporary and historical resequenced genomes from as early as 1829, to track the long-term genomic erosion and immunogenetic diversity decline in this species. 62% of genomic diversity was lost between historical (mean autosomal heterozygosity = 0.00149 ± 0.000699 SD) and contemporary (0.00057 ± 0.000026) parrots. A greater number and length of runs of homozygosity in contemporary samples was also observed. A temporal reduction of the number of alleles at Toll-like receptor genes was found (historical average alleles = 5.78 ± 2.73; contemporary = 3.89 ± 2.10), potentially exacerbating disease susceptibility in the contemporary population. Of particular concern is the new threat of avian influenza strain (HPAI) to Australia. We discuss the conservation implications of our findings and propose that hybridization and synthetic biology may be required to address the catastrophic loss of genetic diversity that has occurred this species in order to prevent extinction.

**Significance statement:** Orange-bellied parrots (*Neophema chrysogaster*) face a dire future, with extinction predicted by 2038 due to severe genetic diversity loss. This Critically Endangered species, now reduced to a single wild population and a captive breeding program, has lost 62% of its genomic diversity since 1829. Contemporary samples show a decline in immunogenetic diversity and signs of very recent inbreeding. Meaning birds today are more susceptible to disease events than birds a hundred years ago. Conservation efforts must consider hybridization and synthetic biology to counteract the catastrophic loss of genetic diversity to ensure the species’ survival. Our study underscores the urgent need for innovative strategies to preserve the adaptive potential and resilience of the orange-bellied parrot and other species in similar situations.

## Introduction

The ongoing climate crisis and ecological destruction are not only impacting human populations-through cropping losses, heat-related deaths and expanding ranges of infectious diseases, but are also placing the worlds’ biodiversity at risk (1–3). At least 32% of known vertebrate species are declining in population size, and 515 vertebrate species have less than 1000 individuals remaining (4, 5). Current conservation efforts range from low intensity species monitoring to intensive *ex-situ* programs, where individuals are bred for release to the wild (for examples see (6–9)). These conservation breeding programs aim to maintain 90% of wild-caught genetic diversity for 100 years and minimise genetic adaptation to captivity (10). However, many *ex-situ* programs often begin with a few founding individuals, making it difficult to preserve genetic diversity long-term, and do not guarantee population recovery (11–13).

Despite conservation efforts, many programs only intervene once significant genetic diversity has already been lost. Intense population bottlenecks incur a genetic drift debt (14) that continues to impact populations even after demographic recovery (15), as small population sizes exacerbate the effects of genetic drift (10). This can lead to increased inbreeding and genomic erosion, reducing adaptive potential, increasing vulnerability to diseases and inbreeding depression, ultimately leading to higher extinction risk (16, 17). Without sufficient baseline data on historical genetic variation, it is difficult to assess the full extent of this loss or to effectively measure the success of recovery efforts (18). Establishing these baselines and understanding the temporal dynamics of genomic erosion are critical to refining conservation strategies, ensuring that actions taken not only focus on population growth but also prioritize the preservation of genetic diversity to maintain long-term adaptive potential (19).

The orange-bellied parrot (*Neophema chrysogaster*) is listed as a Critically Endangered species following significant population declines, which have left it increasingly vulnerable to disease. Historically, the orange-bellied parrot had breeding sites located across the western and southern coasts of Tasmania and migrated in April to their overwinter feeding sites that extended from Adelaide along the southern and eastern coast of Australia to as far north as Sydney (Figure 1; (20)). The species is now restricted to a single breeding site at Melaleuca, SW Tasmania, and its overwinter range is from east of the Murray River in South Australia and west of Port Phillip Bay in Melbourne (Figure 1; (21)). The principal reasons for the population declines through the 20^th^ century are unclear, but are likely due to habitat degradation, with the most recent crash seeing only three wild-born females and 13 males at Melaleuca in 2016 (22–24). Despite regular releases of juveniles from the captive population, the wild population is still small with 81 individuals returning at the start of the 2023/24 breeding season (25). Thus, the species remains impacted by the effects of habitat degradation, inbreeding, low genetic diversity and increased disease susceptibility (21, 26). High juvenile mortality, particularly during migration still presents significant conservation challenges (27). Though inbreeding depression has not yet been studied at the genetic level in this species, evidence from other species with similar demographic trends suggests it likely plays a role in the observed juvenile mortality and reduced fecundity (28, 29).

**Figure 1:**
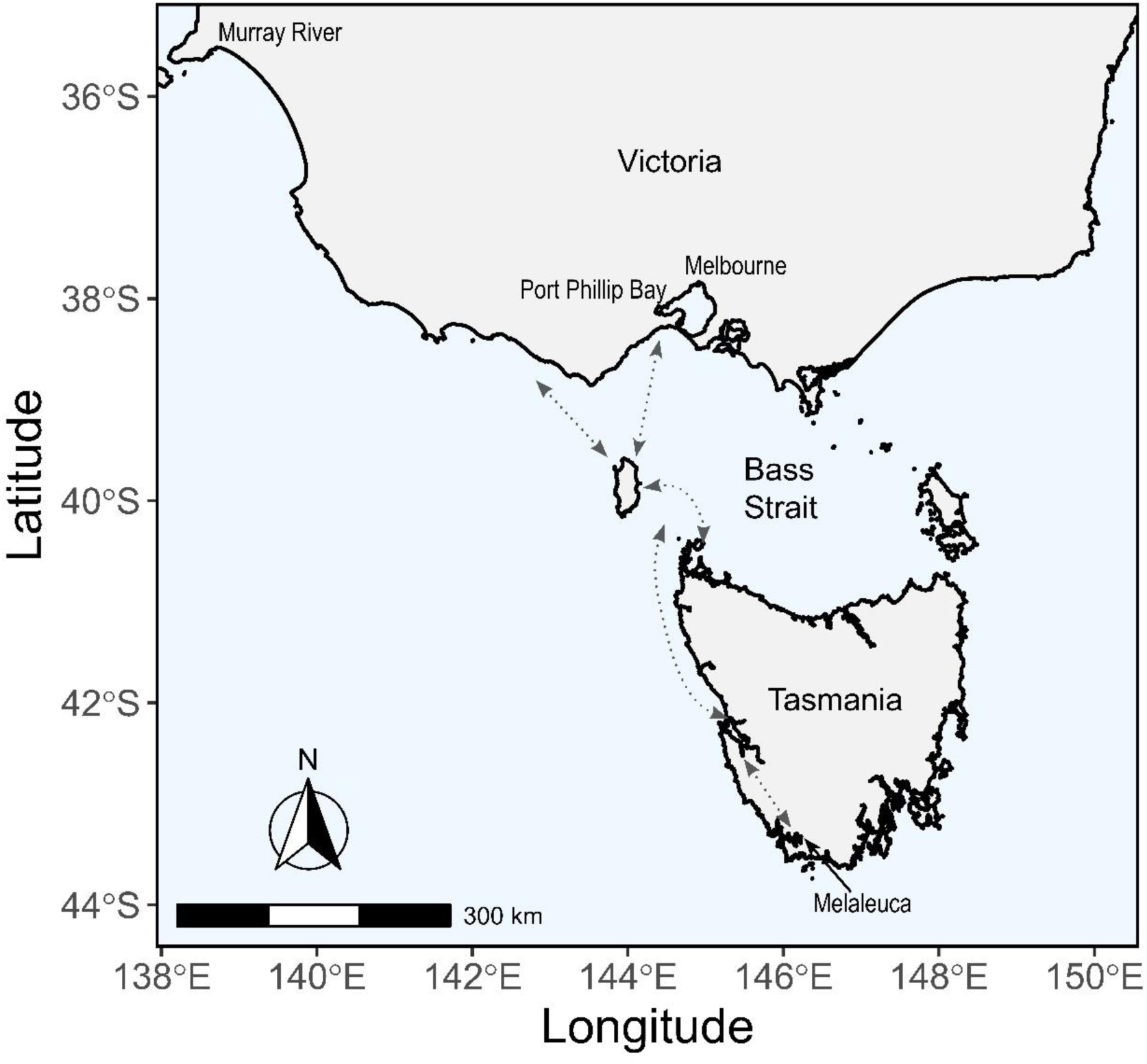
Map depicting the current breeding location at Melaleuca in SW Tasmania, and the over wintering regions in Victoria. The dotted arrows represent the estimated migration route for orange-bellied parrots, but the exact migration flight path is currently unknown. Map modified from Menkhorst, Loyn and Brown (23).

The orange-bellied parrot has been the focus of conservation management since 1979 (22, 30) with an insurance population established in 1986 with an initial intake of ten individuals (22). Although the captive population was supplemented with 44 wild individuals between 1987 and 2011 (31), the current captive population only represents 30 founding individuals and suffers from a number of challenges including disease. Susceptibility to diseases such as beak and feather disease virus (BFDV), Psittacid Adenovirus-2 (PsADV-2) and bacterial infections, *Pseudomonas aeruginosa*, have resulted in mass mortalities in both captive and wild birds in recent years (24, 32–34).

Genetic assessments since 2010 have revealed low genetic variation in wild birds (31, 35) and although the captive population benefited from additional wild bird intakes in 2010, this further exacerbated genetic diversity loss in the wild (31). The species is highly susceptible to genetic component Allee effects, where low genetic diversity impacts fitness (36). Investigations into functional variation using direct sequencing of loci found low (relative to other parrot species) immunogenetic diversity at Toll-like receptors (TLRs), innate immune genes critical to pathogen defence, further underscoring the species’ vulnerability to disease (37).

Due to the continuing loss of genetic diversity, possible inbreeding depression, and risk of stochastic events given the small population size, conservation managers have called for the identification of genetic interventions for the wild population (24). Given that only one wild population remains, extreme measures to introduce genetic diversity have been proposed, such as interspecies hybridisation, using a phylogenetic analysis of the mitochondrial genome to determine the closest relative (38), or to use synthetic biology techniques to re-introduce historical diversity that has been lost via back-breeding, cloning, or genome editing (39). Genetic interventions will require high-quality genetic resources for the species such as a reference genome (40, 41), and an understanding of historic genetic diversity, both of which are lacking for this species. In this study, we aimed to generate the first, high-quality, reference genome of the orange-bellied parrot, and investigate historic and contemporary patterns of genome-wide diversity and immunogenetic diversity.

## Results

### A high-quality reference genome for the orange-bellied parrot

To generate a reference genome for the orange bellied parrot, a total of 18,830 ng of DNA at a concentration of 87 ng/µl was sequenced using a combination of PacBio HiFi and Arima HiC, resulting in 818 Gb of PacBio raw subreads for the *de novo* genome assembly. The genome assembly was assembled into a reference genome of 1.32 Gbp in 2002 scaffolds, with 97.09% of the genome in scaffolds > 50 Kb and a high completeness (>95% complete BUSCOs) (Table 1). HiC scaffolding showed high identity within but not between the eight largest scaffolds (assigned the macrochromosomes), and an expected cluster of microchromosomes as well as several small, unplaced scaffolds (Figure S1).

**Table 1:** Orange-bellied parrot genome assembly metrics.

| Metric | Value |
| --- | --- |
| Genome size (Gbp) | 1.32 |
| No. scaffolds | 2002 |
| No. contigs | 2387 |
| Scaffold N50 (Mbp) | 104.05 |
| Scaffold L50 (Mbp) | 5 |
| Contig N50 (Mbp) | 7.61 |
| Contig L50 (Mbp) | 50 |
| GC content (%) | 42.90 |
| Max scaffold length (Mbp) | 165.58 |
| Max contig length (Mbp) | 47.87 |
| No. scaffolds > 50 Kb | 803 |
| % main genome in scaffolds > 50 Kb | 97.09% |
| BUSCO (aves_odb10, n=8338) | C: 96.8% [S:96.3%, D: 0.5%]<br>F: 0.5%<br>M: 2.7% |
| BUSCO (vertebrata_odb10, n=3354) | C: 96.0% [S: 95.1%, D: 0.9%]<br>F: 1.0%<br>M: 3.0% |

To identify the potential Z chromosome, we used the sequencing covered by the modern resequenced genomes (details below), coverage was stable across the scaffolds in the known male samples, but the known female sample displayed lower coverage at scaffold 5 along with some other samples of unknown sex (Figure S2A). We did not have any known females in the historic resequenced genome dataset (most were unknown sex); differences in coverage were not as clear due to lower average coverage and greater variation in coverage between samples than within the modern genomes (Figure S2B). Alongside the FGENESH++ annotation of the *DMRT3* gene on scaffold 5, we assigned scaffold 5 as the putative Z chromosome.

We identified a circular scaffold of 19,899 bp length representing the mitochondrial genome (Figure S3). The number and order of genes was identical to the previously characterised mitochondrial genome (35). A total of 301 Mbp (22.79%) of the genome was masked as repeat elements. The most common repeat elements were total interspersed repeats (22.62%) followed by long interspersed nuclear elements (LINES; 12.72%) which were comprised mostly of the L3/chicken repeat 1 (CR1) repeat class (Table S1).

### Transcriptome-guided genome annotation and immune gene characterisation

To aid with genome annotation, we sequenced mRNA from eight organs (liver, spleen, kidney, pectoral muscle, heart, testes, tongue, pancreas) and generated a reference guided transcriptome assembly. Mapping rates of the tissue transcriptomes to the repeat-masked genome ranged from 84.60% (testes) to 91.40% (pancreas) (Table S2). Automated genome annotation with FGENESH++ identified 27,046 genes (including non-coding and pseudogenes). A total of 207,171 exons (median of 5 exons per gene, SD = 8.09) and 180,127 introns (median of 4 introns per gene, SD = 8.09) were identified. These values were similar to those identified in the chicken (*Gallus gallus*) (Warren et al., 2017). The global transcriptome contained 91.5% complete BUSCOs and the protein genome annotation contained 83.5% complete BUSCOs (Table S3).

Manual annotation identified nine TLR genes (*TLR1A, TLR1B, TLR2A, TLR2B, TLR3, TLR4, TLR5, TLR7, TLR15*); three cathelicidin genes (*CATHB1, CATHL3, CATHL2*); 11 β-defensin genes; and the MHC Class I gene, UA (Table S4). The cathelicidin and β-defensin genes followed the conserved gene order as described by Cheng*, et al.* (42), including conserved 5’ and 3’ flanking genes, and clustered with other cathelicidins and defensins from species in the Psittaciformes order (Figure S4 and S5). Three paralogs of AvBD3 and two paralogues of AvBD1 were identified in the genome, although the second AvBD1 was pseudogenised (Figure S5 and Table S4). Similar to other Psittaciformes (42), AvBD6/7 and AvBD14 could not be identified in the genome (Figure S5 and Table S4). We were unable to locate a sequence for *TLR21* or any MHC class II genes. To further investigate the class II MHC genes, we developed primers targeting exon 2 and undertook PCR of a number of OBP samples, including the reference individual. Our PCR investigation into MHC II revealed an amplified product for all four primer sets and all six OBPs investigated (Figure S6), additionally, our blast on a *de novo* transcriptome (assembled using default trinity v2.15.1 (43) parameters [data not shown]) identified two MHC class II genes, one alpha chain and one beta chain.

### Loss of genome-wide diversity in contemporary OBP genomes

We performed Illumina whole genome resequencing of 19 contemporary (collected between 2014-2016) and 16 historical (collected between 1829-1986) OBP samples (Table S5). Contemporary resequenced genomes had high alignment rates to the OBP genome (99.63±0.14%). As expected for historical samples containing endogenous bird DNA but also DNA from contaminants (e.g. bacteria and fungi), the alignment rates were lower (56.16±11.52%). Overall, there is slightly higher depth of coverage for historical samples (9.36±6.17X) than for contemporary samples (8.89±3.38X), although more even depth was seen in the contemporary samples. With several historical samples with coverages higher than 15X and as high as 25X (Table S5). Overall, sequencing depth and alignment quality was high enough to proceed with analysis of all samples.

We identified significant declines in heterozygosity between historical and contemporary samples (Contemporary: 5.75×10^-4^; Historical: 1.49×10^-3^, t=-5.72, P=2.18×10^-6^) representing a 62% loss of genetic diversity. The declines seen in heterozygosity also hold true when all samples were downsampled to a consistent read depth (Figure S7). Further analysis of the historical samples revealed a consistent decline in autosomal heterozygosity since the date of the earliest samples in this dataset (1840s: Figure 2A). PCA of all samples identified two historical samples as outliers that limited our ability to investigate patterns of population structuring (Figure S8). Although these two samples had lower sequencing depth (0-5x), there were other samples with a similar sequencing depth, so sequencing depth alone does not explain this result. Our PCA excluding these two outlier samples showed a clear distinction between contemporary and historical samples, as expected given the strong genetic drift after the population decline (Figure S9). There were also differences observed between modern captive samples collected from 2016 and wild samples from 2020 (Figure S8e). The historical samples (with outliers excluded) show an additional two individuals as outliers and make it difficult to identify any clear population structuring amongst the historical samples (Figure S9c and f). Our PCA results also hold true when all samples are downsampled to a consistent coverage (Figure S10 and S11).

**Figure 2:**
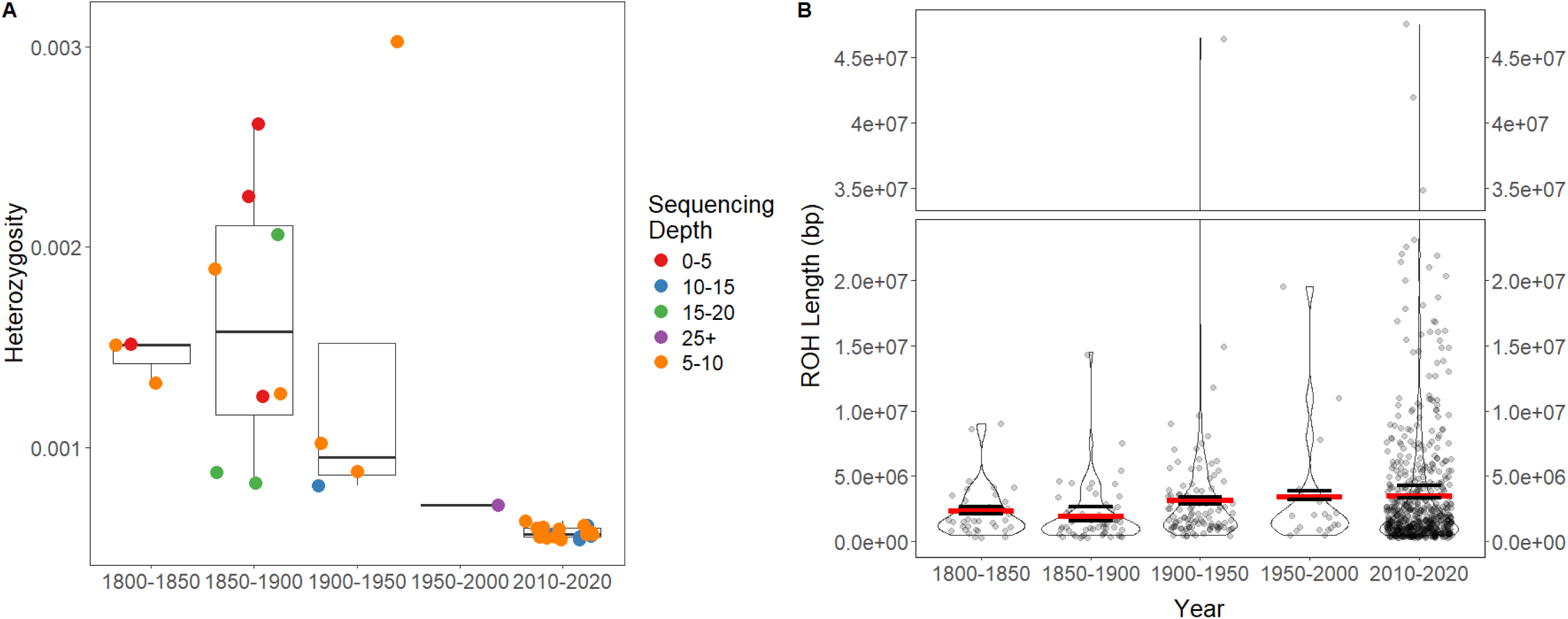
(A) Autosomal heterozygosity as calculated by obtaining the site frequency spectrum using winSFS (v0.7.0). Boxes represent the interquartile range (IQR) with whiskers extending to 1.5*IQR and bold line is the mean for each group. (B) Violin plot of lengths of runs of homozygosity as calculated by ROHan, red line is the average length of ROH as calculated by ROHan and the two black lines represent the minimum and maximum average ROH length.

### Effective population size decline

In order to identify when demographic declines of orange-bellied parrots may have occurred we undertook demographic reconstruction using GoNe (44). Complementary to our genetic diversity results there has been a large decline in the effective population size of OBPs from an estimated Ne of over 1000, 14 generations ago to an estimated Ne of 25-30 in the current generation. Using a generation time of three years (21), we can predict this decline in Ne began around 1978 (Figure 3).

**Figure 3:**
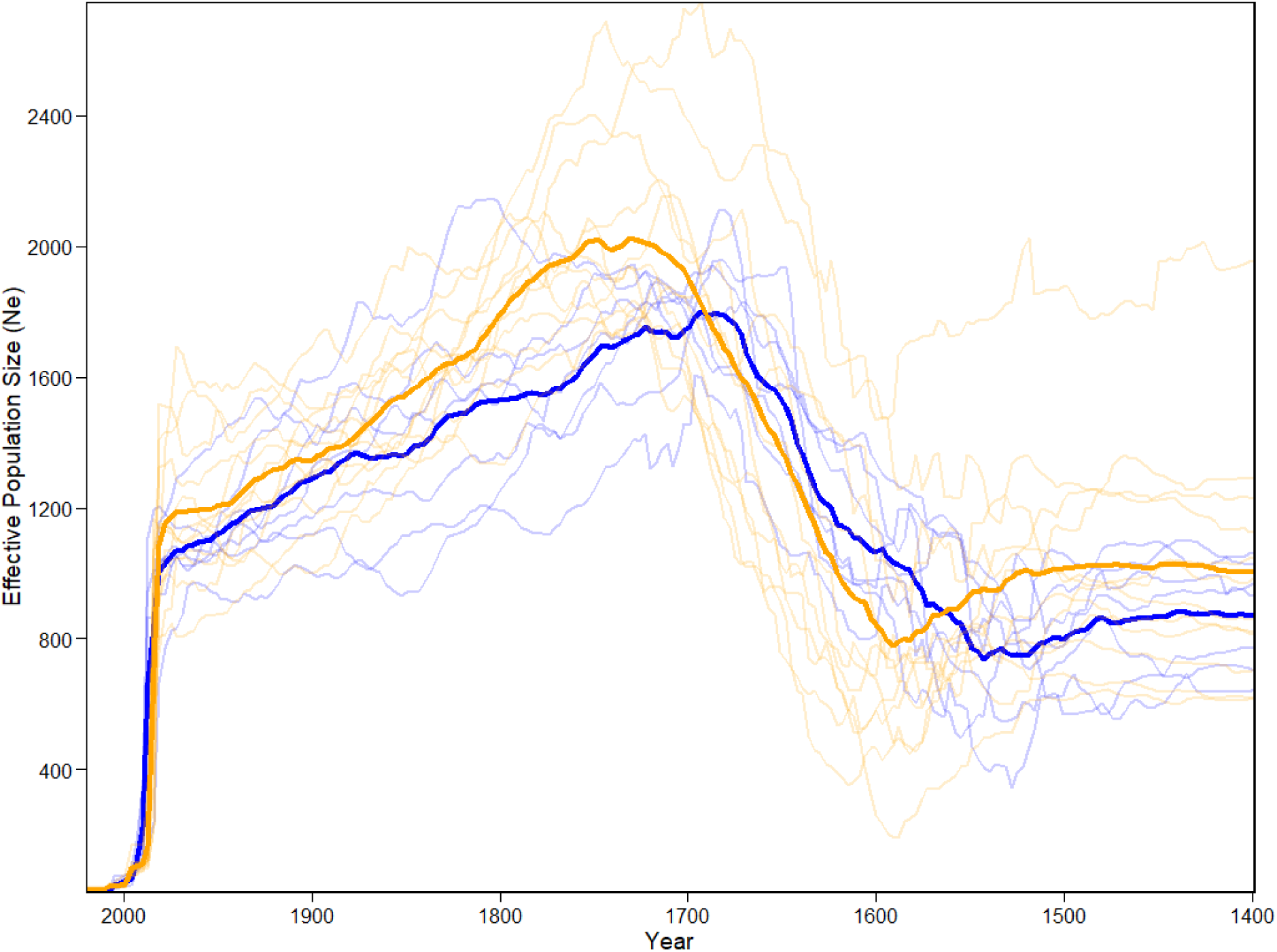
Effective population size (Ne) as calculated by GoNe. Blue lines represent data calculated from birds collected in 2016 and orange lines birds collected in 2020. Faded lines are results of each jack-knifed run of GoNe and solid lines are the average across all jack-knifed run. The year on the x axis was calculated by using a generation time of 3 years, with results for the previous 200 generations shown.

To investigate how inbreeding has changed over time we identified runs of homozygosity (ROH) across contemporary and historical samples using ROHan (45). The contemporary samples had a greater proportion of the genome in ROH >500Kb than the historical samples (Contemporary = 13.42% ± 4.64 SD; Historical = 6.44% ± 4.28 SD; Table S6). ROH were distributed across the autosomal macrochromosome scaffolds (Figure 4). As well as having more ROH than historical samples, the ROH in the contemporary samples were longer (Modern = 3.50×10^6^±1.07×10^6^; Historic = 2.52×10^6^±9.61×10^5^; Figure 2B). We also identified a significant increase in F_ROH 500Kb_ in contemporary individuals compared to historic samples (Modern = 0.129±0.044 SD; Historic = 0.062±0.040 SD; t=4.118; p=2.90×10^-4^). To investigate potential functional impacts of regions of the genome with a high proportion of ROH we located 135 genes covered by ROH in seven individuals. A GO analysis clustered these 135 genes into 38 GO terms (Table S7), visualisation of these GO terms in semantic space with Revigo clustered gene terms in biological processes including, embryo development, anatomical structure development and immune system processes (Figure S12).

**Figure 4:**
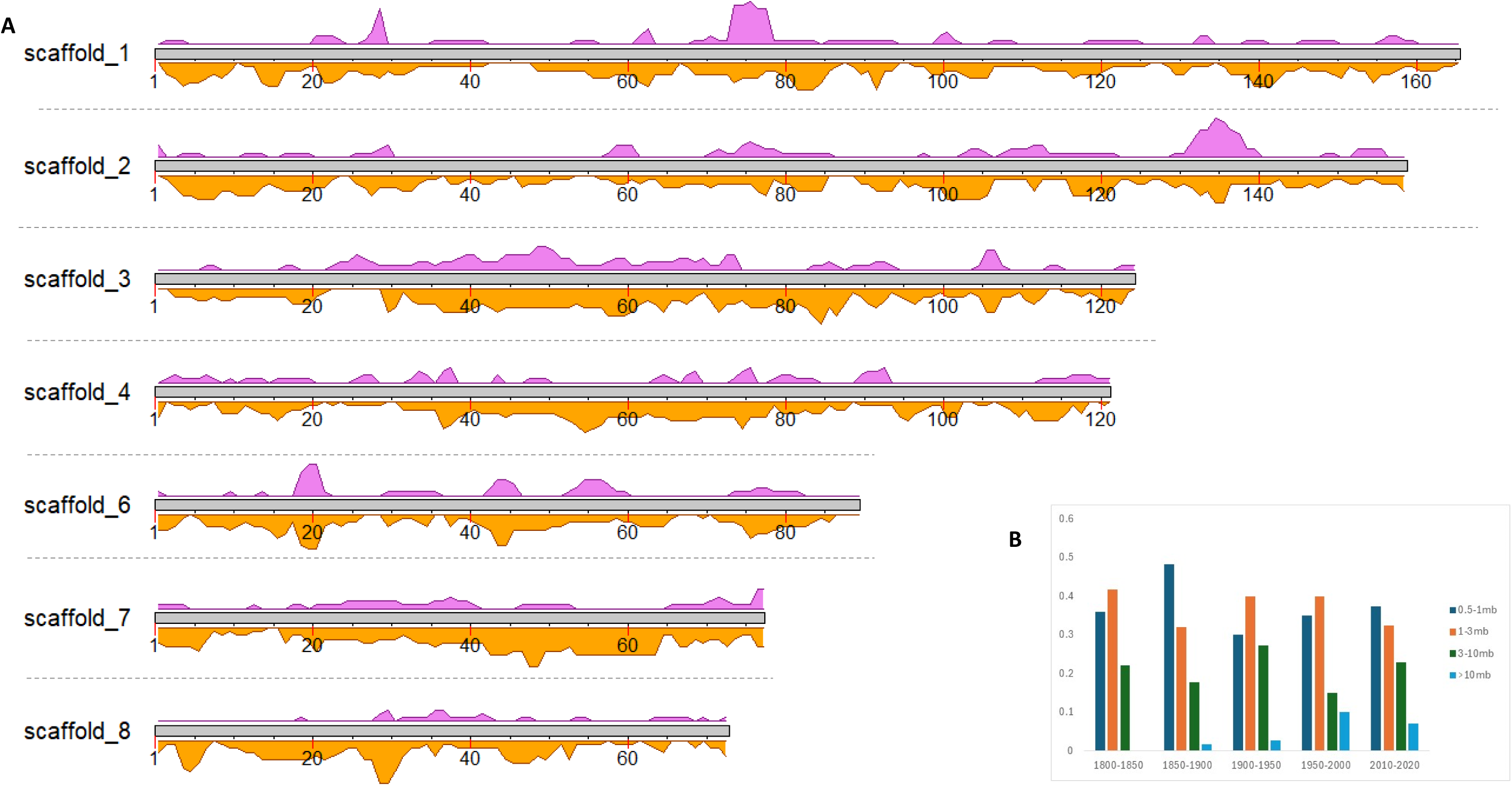
(A) Runs of homozygosity (ROH) in the contemporary (n=19) and historical (n=12, four removed as they did not have the required 6x coverage (45)) resequenced genomes per macro chromosome (excluding sex scaffold). Each chromosome is separated by a grey dotted line. The lilac (historical) and orange (contemporary) density plots show the density of ROH across the entire genome (i.e. higher peaks represent more ROH in a region) Note only scaffolds representing macrochromosomes were included in the analysis and scaffold 5 was excluded as the putative Z sex chromosome. (B) The percentage of the genome in each ROH size class over time.

### Changes in *TLR* diversity over time

We used the resequenced genomes to assess changes in diversity in the toll-like receptor (TLR) genes as extremely low diversity has been previously identified in OBP TLR genes and there have been many associations made between variants in TLR genes and disease outcomes in birds (37, 46, 47). After filtering we identified 46 SNPs in TLR genes, with 20 non-synonymous positions (Table 2). *TLR15* was the most variable gene with nine SNPs, of which six were non-synonymous. *TLR5* was the least variable with a single SNP that resulted in a premature stop codon, although this variant was only present in three individuals in the historical group, all in the heterozygous state. Phasing resulted in 54 alleles, with *TLR15* the most variable with 13 alleles (Table 2, Figure S13). For all TLR genes, we identified fewer numbers of alleles in the contemporary samples compared to the historical samples with the largest decreases seen in *TLR7* (six alleles in the historical samples and two in the modern samples) and *TLR15* (12 in the historical samples and eight in the modern samples) (Table 2). Allelic diversity generally declined from historical to contemporary samples with reductions seen in *TLR1B, TLR2B, TLR3, TLR4, TLR5, TLR7* and *TLR9* (Table 2). Our selection tests failed to identify any sites under selection with the MEME test of selection, however, we identified 9 codons under selection using FEL (Table S8). None of these codons were impacted by an amino acid change.

**Table 2:**
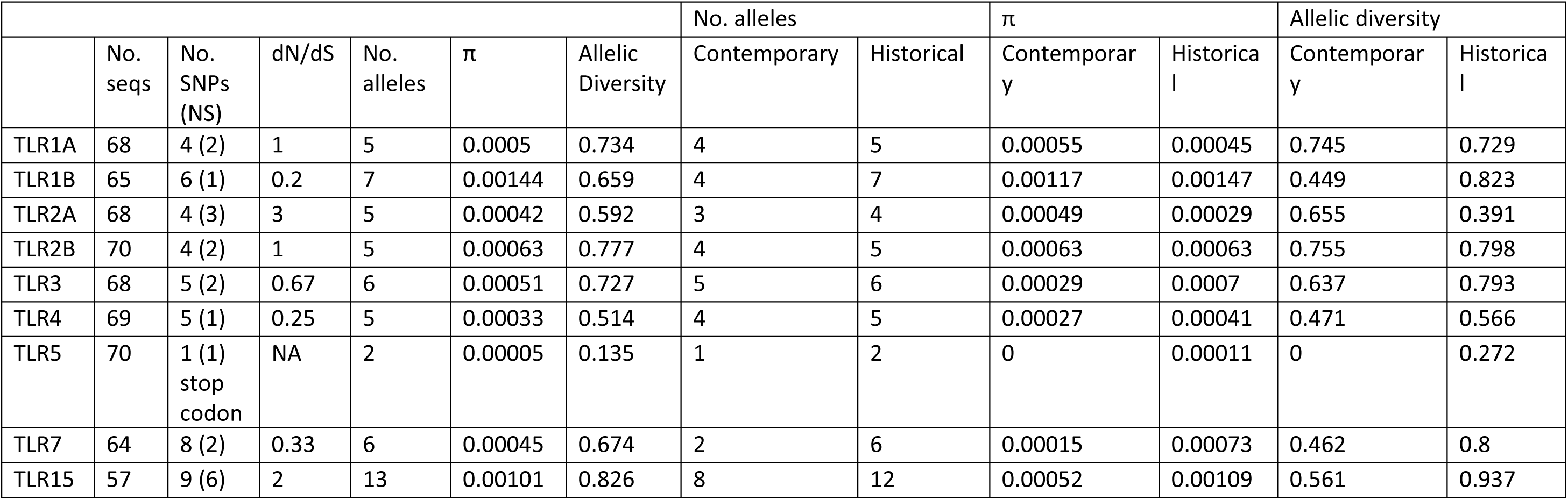
Summary statistics of TLR alleles as generated by DNAsp. NS: non-synonymous.

| | | | | | | | No. alleles | | $\pi$ | | Allelic diversity | |
| --- | --- | --- | --- | --- | --- | --- | --- | --- | --- | --- | --- | --- |
| | No. seqs | No. SNPs (NS) | dN/dS | No. alleles | $\pi$ | Allelic Diversity | Contemporary | Historical | Contemporary | Historical | Contemporary | Historical |
| TLR1A | 68 | 4 (2) | 1 | 5 | 0.0005 | 0.734 | 4 | 5 | 0.00055 | 0.00045 | 0.745 | 0.729 |
| TLR1B | 65 | 6 (1) | 0.2 | 7 | 0.00144 | 0.659 | 4 | 7 | 0.00117 | 0.00147 | 0.449 | 0.823 |
| TLR2A | 68 | 4 (3) | 3 | 5 | 0.00042 | 0.592 | 3 | 4 | 0.00049 | 0.00029 | 0.655 | 0.391 |
| TLR2B | 70 | 4 (2) | 1 | 5 | 0.00063 | 0.777 | 4 | 5 | 0.00063 | 0.00063 | 0.755 | 0.798 |
| TLR3 | 68 | 5 (2) | 0.67 | 6 | 0.00051 | 0.727 | 5 | 6 | 0.00029 | 0.0007 | 0.637 | 0.793 |
| TLR4 | 69 | 5 (1) | 0.25 | 5 | 0.00033 | 0.514 | 4 | 5 | 0.00027 | 0.00041 | 0.471 | 0.566 |
| TLR5 | 70 | 1 (1) stop codon | NA | 2 | 0.00005 | 0.135 | 1 | 2 | 0 | 0.00011 | 0 | 0.272 |
| TLR7 | 64 | 8 (2) | 0.33 | 6 | 0.00045 | 0.674 | 2 | 6 | 0.00015 | 0.00073 | 0.462 | 0.8 |
| TLR15 | 57 | 9 (6) | 2 | 13 | 0.00101 | 0.826 | 8 | 12 | 0.00052 | 0.00109 | 0.561 | 0.937 |

## Discussion

The orange-bellied parrot is listed as one of Australia’s most critically endangered bird species and is predicted to go extinct as early as 2038. Here we have generated high-quality genomic resources for this species, enabling the investigation of changes in the genomic diversity in this species between historic (1800-1900s) and contemporary (2010s) samples. We show a 62% decline in autosomal heterozygosity and increased inbreeding over 200 years. Notably, genetic diversity has also been lost at innate immune genes, including the toll-like receptors, which may explain the species’ susceptibility to recent regular disease events. This loss of genomic diversity threatens the species’ long-term adaptive potential, underscoring the urgent need for informed genetic management to support its recovery.

### Genomic diversity loss and increased inbreeding suggest genomic erosion

We have presented multiple lines of evidence of significant decline in the genomic health of OBPs over the past 200 years. This is noted from the strong reduction over time in autosomal heterozygosity and the marked decline in effective population size during the late 20^th^ century in line with known population bottlenecks (23). Additionally, the increased percentage of genome and average length of ROH in contemporary samples are indicative of recent inbreeding (48). Further evidence of recent inbreeding can be seen by the increase in F_ROH_ in the contemporary samples compared to the historical samples. In samples collected prior to 1900, 42% of the ROH were between 500kb and 1Mb, indicating these ROH first occurred in the genome between 25 and 50 generations ago (75-150 years, using a generation time of three years), with less than 1% of the ROH greater than 10Mb (Figure 4B). This suggests that the ancestral OBP population had some historic background inbreeding, either because of their long-term small population size (see GONE result) or because of an ancestral bottleneck (before 1700’s). Contrary, 7% of ROH in the contemporary samples are greater than 10Mb, indicating these ROH are the result of very recent inbreeding occurring around 2-3 generations (6-9 years) ago. The percentage of the genome in ROH (13.43%) for contemporary, orange-bellied parrots is slightly less but comparable to another critically endangered parrot species with genomic resources, the kākāpō (*Strigops habroptilus*). The extinct historical mainland population of kakapo had 15% of its genome in ROH and the highly inbred extant population on Stewart Island has 53% of their genome in ROH (49). Although ROH were generally spread across the genome, we identified several regions containing a high density of ROH (Figure 4A), this indicates there may be numerous coding regions of the genome which OBPs have little to no variation. Our GO analysis did not identify a single enriched pathway in genes covered by many ROHs, but instead genes involved in several biological processes including, embryo development, anatomical structure development and immune system processes, all of which are known issues to impact OBP viability (24), suggesting that inbreeding depression (if present) is driven by loss of genome-wide variation (Figure S7).

The high disease susceptibility and low juvenile survival of OBPs, could be signs of reduced fitness because of inbreeding depression (50, 51). For example, Kardos*, et al.* (51) identified a 41% lower lifetime reproductive success in inbred killer whales compared to non-inbred counterparts. Our finding of increased F_ROH_ in the contemporary samples compared to historical samples indicates that the current contemporary population suffers from inbreeding. This suggests that intense genomic erosion due to inbreeding could diminish the long-term viability and heighten the extinction risk of the orange-bellied parrot (16). Analysis of individuals with known lifetime fecundity and disease response data would allow the identification of any functional impacts of genomic erosion and inform further whether or not there is a genetic component to some of the observed breeding and disease issues in both the captive and wild populations (52, 53).

### Developing valuable genomic resources to extract challenging immunogenetic information

The genomic resources generated here are the first for the *Neophema* genus contributing to the ever-growing data repository for investigating global bird diversity (54). This high-quality genome and associated global transcriptome data, has permitted the investigation of the immune gene region for this critically endangered species. Manual annotation of immune genes is essential for investigating immune function due to the inability of automated systems to annotate these genes well (55). The TLR gene family are part of the innate immune system and are known to play a fundamental role in pathogen recognition (56). Overall, we found similar variation in OBP TLR genes as another study utilising WGS of a threatened bird, the tchūriwat’/tūturuatu/shore plover (*Thinornis novaeseelandiae*) (57). Previous work using primers for the TLR regions and samples from 20 orange-bellied parrot captive individuals collected in 2013 and 2015, noted that the TLR3, TLR5 and TLR7 fragments were monomorphic, whilst TLR1A and B, TLR10 and TLR4 had low polymorphism (37). Here we sequenced the complete TLR region in 19 contemporary samples and 16 historic samples. The number of alleles observed in our contemporary samples is greater than previously reported, particularly for TLR3, highlighting the higher resolution of genomic resources. Notably, contemporary sample allelic diversity is lower than the historic samples for TLR1B, TLR3, TLR4, TLR5, TLR7 and TLR15. This low diversity potentially explains the ongoing issues with disease outbreaks observed in this species. Further, the contemporary individuals in our study are also monomorphic at TLR5. This is important as TLR5 has been specifically linked to immune recognition of *P. aeruginosa* in mammals (58, 59) and reptiles (60). It has been previously theorised, that low variation at the TLR3 genes may explain the ongoing susceptibility of orange-bellied parrots to PBFV (37) as there is a tentative link between PBFD immunity/susceptibility and TLR3 diversity in red-crowned parakeets (*Cyanoramphus novaezelandiae*) (61). However, as there is little variation between the historical and contemporary samples at the TLR3 gene, and evidence that suggests PBFD has been occurring in wild Australian birds for more than 120 years (33), it makes sense that the historical variation is maintained as infectious diseases are a strong selective pressure that can play a role in immune gene evolution (62). Of particular concern, however, is the new threat of avian influenza strain (HPAI, clade 2.3.4.4b) to Australia that has caused a large number of deaths in a range of bird and mammal species across the globe (63). It has been shown that HPAI viruses cause activation in TLR7 in geese, and are responsible for activation in TLR 1, 2, 4, 5 and 15 in other species infected with avian influenza (64). What this means for the orange-bellied parrot in the longer term is unknown, but limited diversity at the TLR gene region will likely be problematic.

Despite generating a scaffolded long-read genome assembly we were unable to identify any MHC class II genes within the assembly. There are noted difficulties in analysing MHC genes in bird genomes, as they are highly repetitive and have a high GC% (65, 66). The first chicken genome only identified a 92kb region containing 19 MHC related genes and proposed that birds contain a ‘minimal essential MHC’ (67), however, an improved chicken genome assembled with long reads has extended the MHC (68). Despite being unable to identify the class II region in our genome, we did identify a single A and B gene in the *de novo* transcriptome that is consistent with findings of other *Psittaciformes* species (69). It is hoped that by combining our current assembly with future ONT sequencing data, we may be able to resolve the MHC region of the orange-bellied parrot.

### Sustained genomic erosion has worrying implications for conservation and management

Our work highlights the importance of genomic resources and museum collections that allow for long term temporal investigations (19). By utilising museum collections, we have identified higher levels of genome-wide historical diversity, in addition to diversity in key immune genes that have now been lost from this species (22, 23). Our work fits into a broader discussion regarding the best management techniques to retain as much genetic diversity as possible in highly managed populations (70, 71). The orange-bellied parrot has already incurred an important loss of genetic diversity that will likely be compounded by ongoing and future genomic erosion due to the drift debt (14), where continued depletion of genetic diversity is predicted, even after successful demographic recovery (15, 72). Therefore, at this time there are limited options for how to retrieve diversity that has already been lost, as waiting for diversity to be regained through naturally occurring mutations is far too slow, upwards of 100s of generations often requiring extremely large demographic populations (73–75). Current management strategies have to date prevented the extinction of the orange-bellied parrot (12), yet there is still an 87% likelihood they will become extinct in the next 20 years. Our work suggests that now is the time to intervene and reintroduce genetic diversity before it is too late (38, 41).

One potential option is hybridisation with another *Neophema* species (38) as this approach may introduce new genes or genetic variation that increases the adaptive potential of the orange-bellied parrot (76). Another strategy is to reintroduce variation that was previously present in historical populations of orange-bellied parrots through synthetic biology methods (41). Gene editing techniques, like clustered regularly interspaced short palindromic repeats (CRISPR-Cas9) technology have made significant strides in chicken genetics (77), and could be used to mitigate direct population impacts in other bird species, including disease susceptibility (78). Due to the critical nature of the state of orange-bellied parrot genetic diversity, we would recommend investment into both approaches, hybridisation and biotechnology, as both still require further work and development. A two-pronged approach would ensure that time is not wasted on one approach over the other, i.e. 1) future work utilising reference genomes and comparative studies of other *Neophema* species should extend on the work by Hogg*, et al.* (38) and identify the best species to hybridise with; and 2) undertaking further analysis of orange-bellied parrots with known lifetime fecundity and disease responses that will allow us to identify genes involved in these important biological processes so we can use synthetic biology techniques to reintroduce variants identified in historical genomes. Although we note that these approaches must be carefully considered within an ethical framework and require the social licence (38, 79, 80), we also advocate for not delaying further as we need to solutions to improving genetic diversity in this critically endangered species.

Here we have provided high-quality genomic resources for the orange-bellied parrot and show that we have reached a tipping point – the species will go extinct due to genetic erosion unless we intervene. Are we prepared to resurrect genetic diversity that was lost over the past 200 years? Or are we prepared to hybridise them with another species? The basic question is, are we prepared to use genomic approaches to bring a species back from extinction? If we wait too long it will be too late.

## Materials and methods

### Reference genome and transcriptome sampling, extraction, and sequencing

Two captive adult male OBPs were euthanised on 20/08/2020 for medical reasons and immediately dissected for tissue collection. Tissues were collected in duplicate for DNA sequencing (flash frozen at −80°C) and RNA sequencing (fixed in RNALater for 24 hours before freezing at −80°C) under NSW Scientific Licence SL101204. High-molecular weight DNA was extracted from the heart, kidney and pectoral muscle tissue of one bird (SB# 2035) using the Circulomics Nanobind Tissue Big DNA Kit v1.0 11/19. DNA quality and concentration was assessed by Nanodrop (ThermoFisher Scientific) and Qubit fluorometer dsDNA BR assay kit (ThermoFisher Scientific). DNA from the three tissues was pooled for sequencing. PacBio HiFi library preparation and sequencing was performed at the Australian Genome Research Facility (AGRF) using 1 SMRT cell on Sequel II. Heart tissue flash frozen at −80C was sent for HiC library preparation using the Arima HiC kit and sequencing on an Illumina NovaSeq 6000 as 150 bp paired end reads at the ACRF Biomolecular Resource Facility (The John Curtin School of Medical Research, Australian National University, Canberra, Australia).

For transcriptome sequencing, RNA was extracted from tissue for both birds using the Qiagen RNeasy Mini Kit with DNAse treatment. RNA quality was assessed by Nanodrop and BioAnalyzer to obtain RIN scores. The liver, spleen, kidney and pectoral muscle of the reference genome bird, and the heart, testis, tongue and pancreas of the second bird underwent library preparation (Illumina Stranded mRNA Prep Ligation) and 100-bp paired-end sequencing on a NovaSeq 6000 S1 flowcell at Ramaciotti Centre for Genomics (University of New South Wales, Sydney, Australia).

### Resequenced genome sampling, extraction, and sequencing

We generated resequenced genomes for 19 contemporary orange-bellied parrots (OBPs) (collected between 2014 and 2016 from the captive population (Taroona; N=7), or in 2020 from the wild population at Melaleuca; N=12) and 16 historic samples that were collected since 1843 from South Australia and Victoria and obtained from natural history collections (Table S5). Contemporary samples were obtained from the OBP biobank at the Australian Museum Research Institute. DNA was extracted from contemporary samples from dried blood on FTA or filter paper using the DNeasy Blood and Tissue Kit (Qiagen, Hilden, Germany) with the protocol for nucleated blood and submitted to BGI-Europe for library preparation and paired-end 150 sequencing on the MGI-Tech DNBSEQ-G400. Historical toe-pad samples were obtained from the South Australia Museum, Museums Victoria (Australia) and the Muséum National d’Histoire Naturelle (France). Historical samples were extracted following Gilbert, Moore, Melchior and Worobey (81) in ultra-clean laboratories for ancient DNA at the University of Copenhagen. Genomic libraries were built following Kapp, Green and Shapiro (82) as modified for the BGISEQ sequencing platform (83). Indexed libraries were amplified for 20 PCR cycles and submitted to BGI-Europe for paired-end 100 sequencing in their DNBSEQ-G400.

### Genome and mitogenome assembly

PacBio reads were converted to FASTQ format using SAMtools v1.11 bam2fq (84). Q20+ HiFi reads were retained using BamTools v2.5.1 filter “rq”:”>=0.99” (https://github.com/pezmaster31/bamtools). HiFi reads with adapter contamination were identified and removed using HiFiAdapterFilt with default parameters (85). *De novo* genome assembly was performed using hifiasm v0.16.1-r375 (86) with the parameters-a 6 (six rounds of assembly graph cleaning) and -s 0.65 (similarity threshold for purging of duplicate haplotigs) on a Pawsey Supercomputing Service Nimbus virtual machine (64 vCPUs; 256 GB RAM; 3 TB storage). HiC reads were aligned to the assembly following the Arima Genomics Mapping Pipeline (A160156 v2; https://github.com/ArimaGenomics/mapping_pipeline) and scaffolded with YaHS (87). The HiC contact matrix was generated with Juicer v2.17 and visualised with JuiceBox (88). Genome assembly statistics were calculated with bbmap v38.86 ‘stats.sh’ (https://sourceforge.net/projects/bbmap/). BUSCO v5.5.0 (89) was used to assess assembly completeness with both the vertebrata_odb10 and the aves_odb10 lineage datasets, implemented on Galaxy Australia (90).

The OBP mitochondrial genome was previously characterised using Roche 454 next generation sequencing data (35). In this study, we identified the mitogenome from our long-read genome assembly using MitoHiFi v2 (91), using the original mitogenome NCBI accession number NC_019804.1 as input, and compared the gene order to that of the original. We visualised the mitogenome with MitoZ v2.3 (92).

### Transcriptome assembly

Quality control of raw transcriptome reads was visualised using FastQC v0.11.8 (93) and MultiQC (94) to gather reports. Reads were quality trimmed using trimmomatic v0.39 (95) with ILLUMINACLIP: TruSeq3-PE to remove Illumina adapters, SLIDINGWINDOW:4:5, LEADING:5, TRAILING:5 and MINLEN:25. FastQC was again run to assess quality after trimming. FASTQ files of paired reads for transcriptomes run across multiple lanes were merged. The repeat-masked genome (see below) was indexed for alignment of transcriptome reads to the genome with hisat2 v2.1.0 (96). SAMtools v1.9 (84) was used to convert SAM files to BAM format and sort by coordinate and stringtie v2.1.6 (97) was used to generate a GTF file.

A global transcriptome (consisting of transcripts from all tissues) was created to provide mRNA evidence for genome annotation. Aligned reads were merged into transcripts using TAMA (98) and python v2.7.9 with -m 3 -z 500. Transcripts were filtered to remove transcripts with weak evidence, defined as transcripts found in only one tissue and with fragments per kilobase of transcript per million mapped reads (FPKM) < 0.1. bedtools v2.29.2 (99) was used to create a FASTA file of retained transcripts. Coding prediction was performed using CPC v2.0 (100). Open reading frames ORFs > 100 amino acids long default were extracted and coding regions predicted using TransDecoder v2.0.1 (101). Candidate peptides were screened for homology against the Swissprot protein database (102) using blastp from BLAST+ v2.7.1 (103) with an E-value threshold of 1e-5. BUSCO v5.5.0 (89) was used to assess alignment of the global transcriptome and of individual tissue transcriptomes using transcriptome mode and the aves_odb10 lineage dataset.

### Genome annotation

RepeatModeler v2.0.1 (104) was used to build a *de novo* database and model repeats from the genome assembly. RepeatMasker v4.0.9 (105) was then used to classify repeats and mask them from the unfiltered genome assembly for transcriptome alignment and genome annotation. The - nolow parameter was specified to prevent masking low complexity DNA or simple repeats. The global transcriptome was converted for input to FGENESH++ (106) using samtools v1.9, seqtk v1.3 and seqkit v0.10.1, retaining only the longest complete transcripts with an open reading frame for each predicted gene to use as mRNA evidence. FGENESH++ annotation was undertaken with general non-mammalian parameters, an optimised gene-finding matrix from the American crow (*Corvus brachyrhynchos*) and the mRNA evidence, with the NCBI non-redundant protein database used for homology-based gene predictions. The annotation was assessed using genestats to summarise the number and length of exons and introns and BUSCO v5.5.0 with the aves_odb10 lineage dataset and “proteins” mode.

### Immune gene annotation

As automated annotation pipelines are unreliable at annotating immune genes, especially in non-model organisms (55), we used a homology-based approach via BLAST v2.3.30 (103) and/or a motif-based approach via HMMER v3.3.2 (hmmer.org) to annotate major histocompatibility complex (MHC), Toll-like receptor (TLR), and two families of antimicrobial peptide genes (cathelicidins and β-defensins) in the OBP in the genome.

For antimicrobial peptide genes, we used query sequences collated from 53 avian species by Cheng*, et al.* (42) to search the genome via BLASTn and the global transcriptome using tBLASTn with an e-value threshold of 1e-10. Unique hits with the smallest e-value were selected for further investigation and sequences extracted by bedtools for manual inspection of the open reading frame. To confirm putative cathelicidins, protein sequences were aligned to known cathelicidins from 53 avian species (42) and fish (107) in BioEdit using ClustalW (108). The cathelicidin alignment was used to generate a neighbour-joining phylogenetic tree in MEGA v11 (109) using the p-distance method with pairwise deletion and 500 bootstrap replicates. The defensin alignment was used to generate a maximum likelihood phylogenetic tree in IQ-TREE v1.6.12 (110), with ModelFinder (JTT+G4) (111) and 1000 Ultrafast bootstrap replicates (112).

For TLR genes we used query sequences downloaded from NCBI for chicken (DQ004555.1, DQ267901.1, EF137861.1, FJ915386.1, KF697090.1, KU235484.1, NM_001007488.5, NM_001030558.3, NM_001081709.4, NM_001161650.3, NM_001396826.1) and used blastn with default parameters against the OBP genome.

For MHC genes we used query sequences from Westerdahl*, et al.* (113) and blastn with default parameters against the OBP genome, global transcriptome, and unassembled PacBio HiFi reads. Our blast search with these sequences was unable to identify any sequence matching to MHC Class II genes. We further attempted to identify Class II genes in the OBP by annotating the Class II genes of more closely related Australian parrot species, including the swift parrot (*Lathamus discolor*) (114), western ground parrot (*Pezoporus flaviventris*; GCF_033815535.1) and eastern ground parrot (*Pezoporus wallicus*; GCF_028554395.1). We used these annotated parrot MHC genes to build a HMMER model and ran nhmmer with default parameters against the OBP transcriptome and genome. MHC Class II genes were still unable to be identified and so we developed four sets of primers based on conserved regions of closely related parrot species, exon 2 (Table S9). We performed PCR reactions on *Neophema* and *Neopsephotus* samples from Hogg*, et al.* (38) as well as the reference genome individual (OBP bird SB# 2035) and the swift parrot reference genome individual ((114); Table S10). Primers were mixed in 50µL containing 1x MyTaq Mix (Bio-Line), 1µM of forward and reverse primers, 1µL of DNA (∼50ng/µL) and made up to 50µL with water. PCR conditions on a T100 Thermal Cycler (Bio-Rad) consisted of an initial denaturation step at 95 °C for 1 min followed by 35 cycles of denaturation at 95 °C for 15 s, locus-specific annealing temperature for 15 s and extension at 72 °C for 10 s. Amplification was confirmed on 1% agarose TBE gel stained with SYBR Safe DNA (Life Technologies) alongside Hyperladder IV (Bioline) for 25 min at 90 V. Bands were visualised using a Gel Doc XR + (Bio-Rad) under ultraviolet light and images were analysed with ImageLab (Bio-Rad). The swift parrot was used as a positive control for our PCR as this was one of the species used to design the primer sets, all reactions included a negative control of water. To further investigate if MHC Class II is present, we performed a *de novo* assembly of transcriptomes from bird SB# 2035 using Trinity v2.5.15 (115) on Galaxy Australia (90) and performed blastn as described above with query sequences from Australian parrots.

### Sex chromosome identification

The reference genome was generated from the homogametic sex, a male (ZZ) parrot. To identify the putative sex chromosome, we examined coverage across the eight genome scaffolds representing macrochromosomes using the resequenced genomes to identify hemizygous regions in female (ZW) samples. Coverage was calculated from the reference-aligned resequenced genome BAM files in non-overlapping 1 Mb windows as per Dodge*, et al.* (116) using BEDTools genomecov v2.29.2 (99). In addition, the FGENESH++ automated annotation was searched for the doublesex and mab-3-related transcription factor 3 (*DMRT3*) gene, present on the chicken Z chromosome (117) and implicated in male sex differentiation (NCBI, https://www.ncbi.nlm.nih.gov/gene/58524).

### Resequenced genome alignment and variant calling

Raw fastq reads for contemporary individuals were quality checked and trimmed using fastqc (v0.11.8) and trimmomatic (v0.39) with the parameters ILLUMINACLIP:TruSeq3-PE.fa:2:30:10 SLIDINGWINDOW:4:5 LEADING:5 TRAILING:5 MINLEN:25. Reads were then aligned to the OBP reference genome using the Burrows Wheeler Aligner (BWA; v0.7.17 (118)) with the bwa-mem algorithm and default parameters. Resulting alignment files were sorted into bam format using samtools sort (v1.6) as each individual had been sequenced across multiple lanes, bam files for each lane were then merged using samtools merge and duplicates marked and remove using picard (v2.21.9) MarkDuplicates (http://broadinstitute.github.io/picard/).

Reads from historical samples were first merged and collapsed using fastp v0.22.0 (119) then quality checked using FastQC. Merged and unmerged reads from fastp were aligned to the OBP genome using the bwa-aln algorithm with parameters *-n, -o* and *-l*, set to 0.03, 2 and 1024, respectively. The aln algorithm is the recommended BWA algorithm for reads shorter than 70 bp (118) (Li and Durbin 2009) and has shown good performance for short and damaged reads (120). Alignments were converted to sorted bam format using bwa samse (for unmerged read alignments) and bwa sampe (for merged read alignments); resulting bam alignments were then merged for each sample using samtools merge and duplicates marked using picard MarkDuplicates. As historical DNA can be degraded, damage profiles were assessed using mapDamage v2.3.0a0 (121) downsampled to 50,000 reads, with a read length expected of 150bp.

We used two methods for downstream data analysis of resequenced genomes. Firstly, we used angsd v0.914 (122) with the following parameters, -uniqueOnly 1 -remove_bads 1 - only_proper_pairs 1 -skipTriallelic 1 -rmTrans 1 -C 50 -baq 0 -minMapQ 30 -minQ 20 -doCounts 1 - minInd 2 -doMajorMinor 1 -GL 2 -doGlf 2 -doMaf 2, to generate a lists of sites across; all samples, contemporary samples only and historical samples only. The sites generated across all samples were used for heterozygosity and principal component analysis (PCA) analysis containing all samples, the contemporary and historical only lists were used for PCAs of contemporary and historical samples, respectively. We removed the putative sex scaffold (scaffold 5) and removed all transitions from analysis of historical samples to avoid bias from DNA damage. We examined alignment rates to the genome, percentage of endogenous content, and sequencing depth and determined that sequencing quality was acceptable for further analysis of all samples.

### Genome-wide diversity in contemporary and historic samples

Individual-level autosomal heterozygosity was estimated by obtaining the site frequency spectrum using winSFS v0.7.0 (123) on the site allele frequencies calculated above with angsd and determining the proportion of heterozygous positions. We tested for statistically significant differences between modern and historical heterozygosity with a two-sample t-test at a significance threshold of α=0.05. We also examined autosomal heterozygosity over time in the historic samples by estimating the rate of heterozygosity loss by 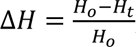, where H is the initial heterozygosity and H_t_ is heterozygosity at time point t. We confirmed out results were not artefacts of sequencing depth differences by downsampling all samples to a 5x depth and repeating the analysis.

We performed a PCA using angsd to output a covariance matrix, with the following parameters, - minMapQ 30 -minQ 20 -GL 2 -doMajorMinor 1 -doMaf 1 -SNP_pval 2e-6 -doIBS 1 -doCounts 1 - doCov 1 -makeMatrix 1 -minMaf 0.05 -P 5, for all samples combined, and historical and contemporary samples separately with PCA visualised using ggplot2 v3.4.4 (124) in R v4.3.2 (125). PCA visualisation of contemporary and historic samples identified two historic samples as outliers, the most distant outlier was the oldest sample in the dataset and was sequenced with the lowest depth and coverage of the genome (42% genome coverage). We repeated the PCA excluding these two samples. We confirmed out results were not artefacts of sequencing depth differences by downsampling all samples to a 5x depth and repeating the analysis.

Runs of homozygosity were identified using all resequenced genomes for the eight macrochromosomes, excluding the sex chromosome (scaffold 5) using ROHan (45) with --rohmu 5e-5, -TsTv 3 and –size 100,000 on an Amazon Web Services ubuntu 20.04 LTS cloud machine (r5.16x large, 64 vCPU, 512 Gb RAM, 1TB attached storage). For the historic samples, the bam2prof function of ROHan was first run to estimate deamination profiles, which were then supplied to the ROH estimation. We removed four historic samples as they had coverage lower than the 6x that is required to reliably calculate ROH of 500kb (OBPI_505, OBPI_508, OBPII_ReHoI_503 and OBPII_ReHoI_505) (45). ROH based inbreeding coefficients (F_ROH_) were calculated based on the proportion of the genome present in ROHs > 500Kb divided by the maximum length of ROH able to be identified across the 7 autosomal macrochromosomes. To further investigate the potential impact of ROHs in contemporary OBP samples we conducted a gene ontology (GO) analysis. To do this we identified which genes (as annotated by FGENESH++) overlapped with ROHs (>500Kb) using bedtools intersect v2.29.2 (99). We retained genes that overlapped with ROHs in at least 7 individuals. We ran blastp of the protein sequences for these 135 genes against the uniport non-redundant protein database, retaining only the top hit for each protein and conducted GO analysis using GoNet (126). The resulting GO terms were clustered and visualized in sematic space using REViGO (127).

### Demographic reconstructions

To investigate historical patterns of demography we calculated effective population size using GoNe (128). We first called SNPs using bcftools v1.17 (84) mpileup with a quality filter of 30 to generate single sample gvcf files. We then used bcftools merge to combine all single sample gvcf into a population level gvcf and bcftools call to retain only variable sites. We performed stringent filtering by retaining variant sites with minor allele frequency > 0.04, minimum average depth of 5, maximum average depth of 200, genotype quality of 10, minimum quality score of 10 and maximum missing data of 10%. As GoNe is impacted by population admixture (44) we ran GoNe on two different groups of individuals for our modern samples as follows; 1 – samples collected in 2020 (N = 12), 2 – samples collected in 2016 (N = 7). For each group we removed any invariant sites, retained only SNPs on the seven autosomal macro-chromosomes and removed transitions from the historical samples. There is a known artefact that occurs when there is population structuring between individuals used in GoNe analysis where in recent generations there is a large increase and sudden drop in Ne, which can be improved through adjustments of the parameter hc, which gives the maximum value of the recombination rate. Therefore, we ran numerous iterations of GoNe to determine the optimal hc (0.01, 0.05, 0.1) and different recombination rate estimates (1, 2, 3). To estimate a confidence interval for our estimates, we performed jack-knifing on the best performing combination of recombination rate (2 cMMb) and hc (0.05), where we iteratively removed one individual at a time and randomly subsampled SNPs in each replicate.

### Functional Analysis of TLR genes

To investigate diversity within the nine annotated TLR genes we performed stringent filtering as described above. To identify full length nucleotide sequences for each individual we phased haplotypes which contained more than two variants using PHASE v2.1.1 (129–131) to identify and assign alleles. We filtered alleles by retaining only those alleles present more than twice in our dataset. For each gene we determined nucleotide and allelic diversity using DNAsp v6.12.03 (132), for all individuals at once as well as modern and historic samples separately. Selection tests were conducted mixed effects model evolution (133) and fixed effects likelihood (134) using HyPHy (135) on the datamonkey server (136) and allele network plots generated using the pegas package v1.3 (137) in R (125).

## Acknowledgements and funding sources

The authors acknowledge the traditional custodians of the land upon which the orange-bellied parrot resides, as well as the custodians of the historic range of the species and pay our respects to their elders past and present. We acknowledge the many years of effort and work of the orange-bellied parrot recovery team in the conservation of this species. We thank S. Troy from the Natural Resources and Environment Tasmania and D. Stojanovic from the Australian National University for collecting contemporary samples and accessioning them to the Australian Museum Research Institute, we thank the curators and staff of the AMRI for providing these contemporary samples. We are grateful to the curators and staff from the South Australia Museum, Museums Victoria, and the Muséum national d’Histoire naturelle for providing historical samples. We thank Priam Psittaculture Centre for sampling of the reference genome individual and transcriptomes. We thank Claudia Fontsere for advice on bioinformatic analyses. The authors wish to acknowledge the use of the services and facilities of the Australian Genome Research Facility; Ramaciotti Centre for Genomics at the University of New South Wales; and the Biomolecular Resource Facility at the John Curtin School of Medical Research, Australian National University. This work was supported by resources provided by the Pawsey Supercomputing Centre with funding from the Australian Government and the Government of Western Australia; the Sydney Informatics Hub, a Core Research Facility of the University of Sydney; Amazon Web Services; RONIN; and Australian Biocommons which is enabled by National Collaborative Research Infrastructure Scheme via Bioplatforms Australia Threatened Species Initiative funding; DNRF143 Center for Evolutionary Hologenomics; ERC CoG 681396 Extinction Genomics; MSCA grant GENDANGERED 840519, and ERC StG 101078303 ERODE. Views and opinions expressed are however those of the authors only and do not necessarily reflect those of the European Union or the European Research Council. Neither the European Union nor the granting authority can be held responsible for them

## Data availability

All the raw data generated for this project is publicly available on NCBI (PRJNA1140271). Additionally, raw data for the genome (HiFi and HiC) and transcriptome are available on the Bioplatforms Australia data portal (https://data.bioplatforms.com/organization/threatened-species). Primers designed to amplify Neophema MHC class II genes are presented in Table S9 and all genomic coordinates for genes annotated are presented in Table S4. All other data is available upon request.

## Project Contributions

Conceptualization – CJH, HEM; Resources – CJH, KB, HEM, TG; Genome and transcriptome sampling – CJH, KAF; Genome extractions – KAF, EP; Transcriptome extractions – KAF, EP; Genome assembly – KAF; Transcriptome assembly – KAF; Resequenced genome data generation – HEM, TG; Resequenced genome analysis – KAF, LS, HEM; MHC – LS, KAF; TLRs – LS; AMPs – EP, KAF; Manuscript original draft – KAF, LS, CJH; Manuscript final draft – KB, HEM, LS, CJH, TG, KAF, EP

---

**Figure S1:**
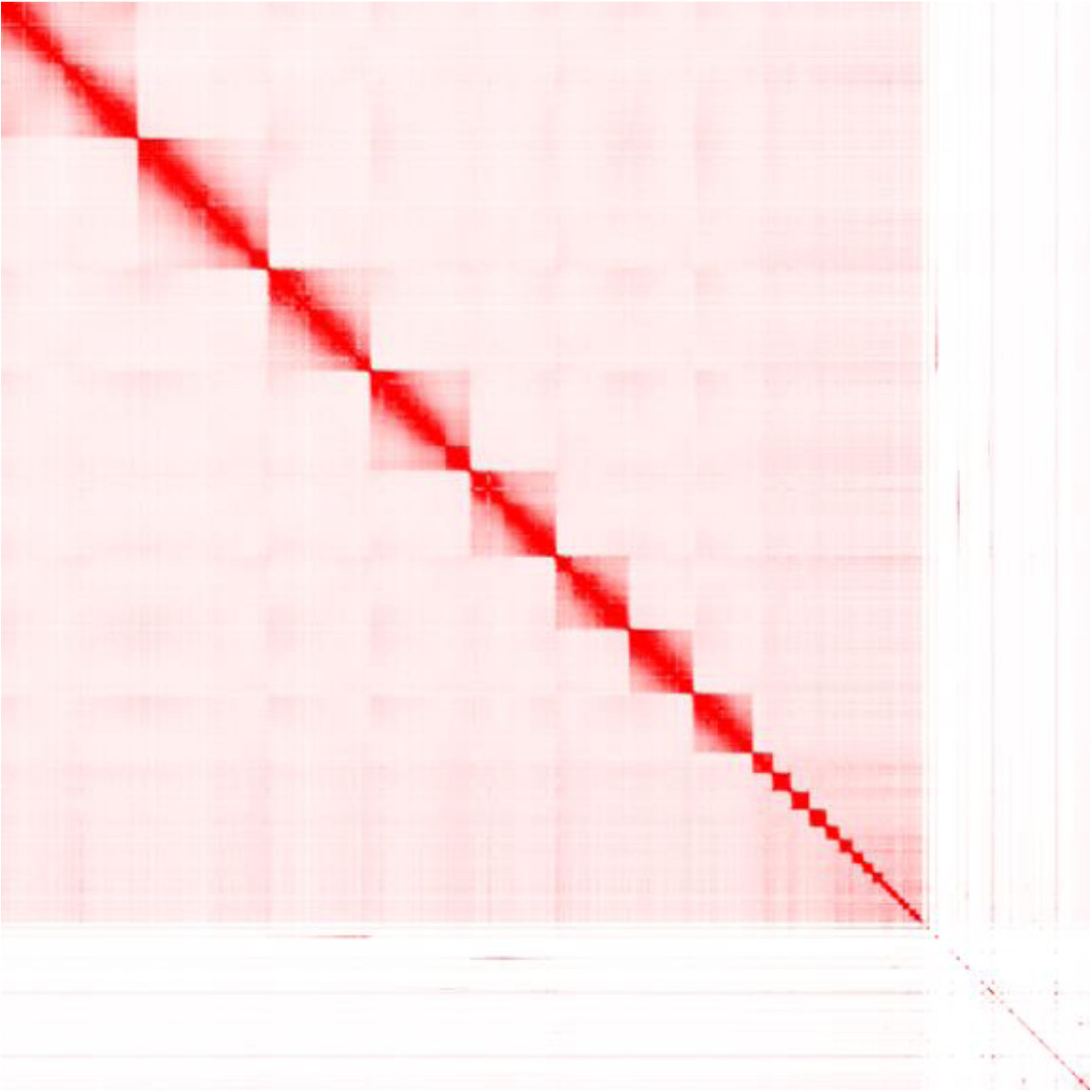
HiC contact map of the scaffolded genome assembly. The map displays the eight macrochromosomes, microchromosome cluster, and unplaced scaffolds.

**Figure S2:**
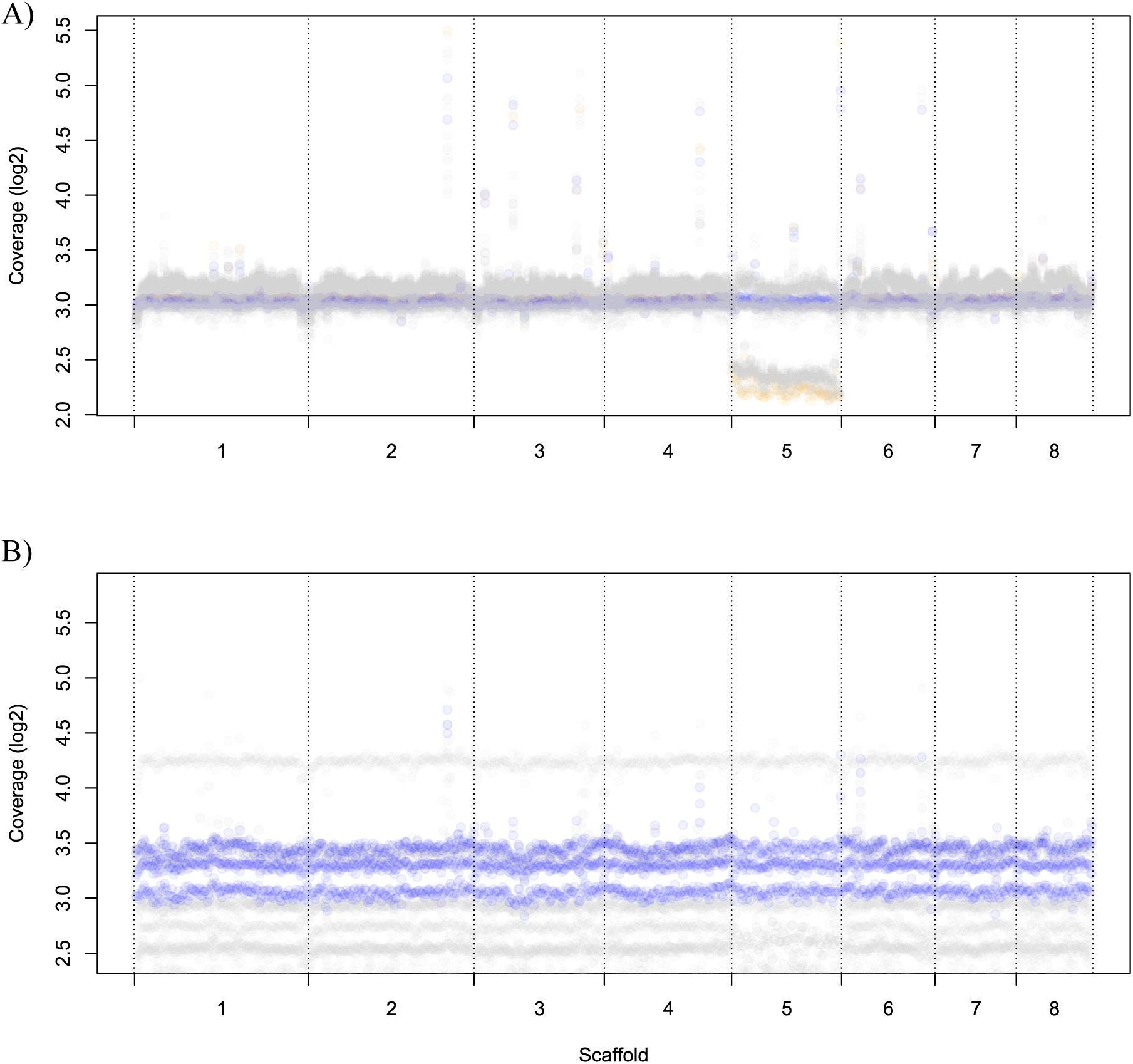
Coverage (log_2_) of the A) modern resequenced genomes and B) historic resequenced genomes, at each 1 Mb window across the eight macrochromosome scaffolds. Points are shaded blue for known males, orange for known females, and grey for unknown sex. Note there were no known females in the historic samples.

**Figure S3:**
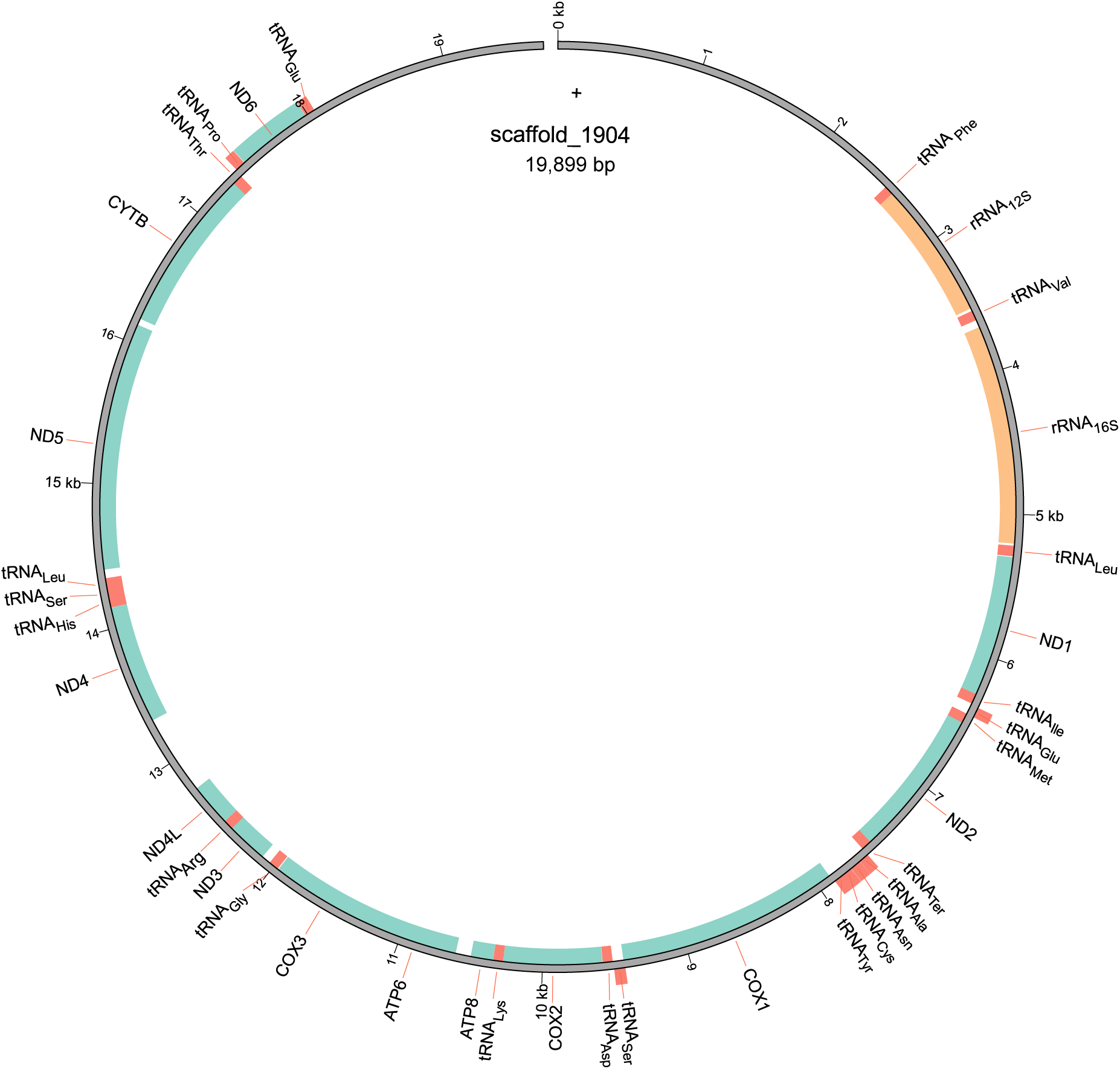
The mitochondrial genome of the orange-bellied parrot (*Neophema chrysogaster*) characterised in this study.

**Figure S4:**
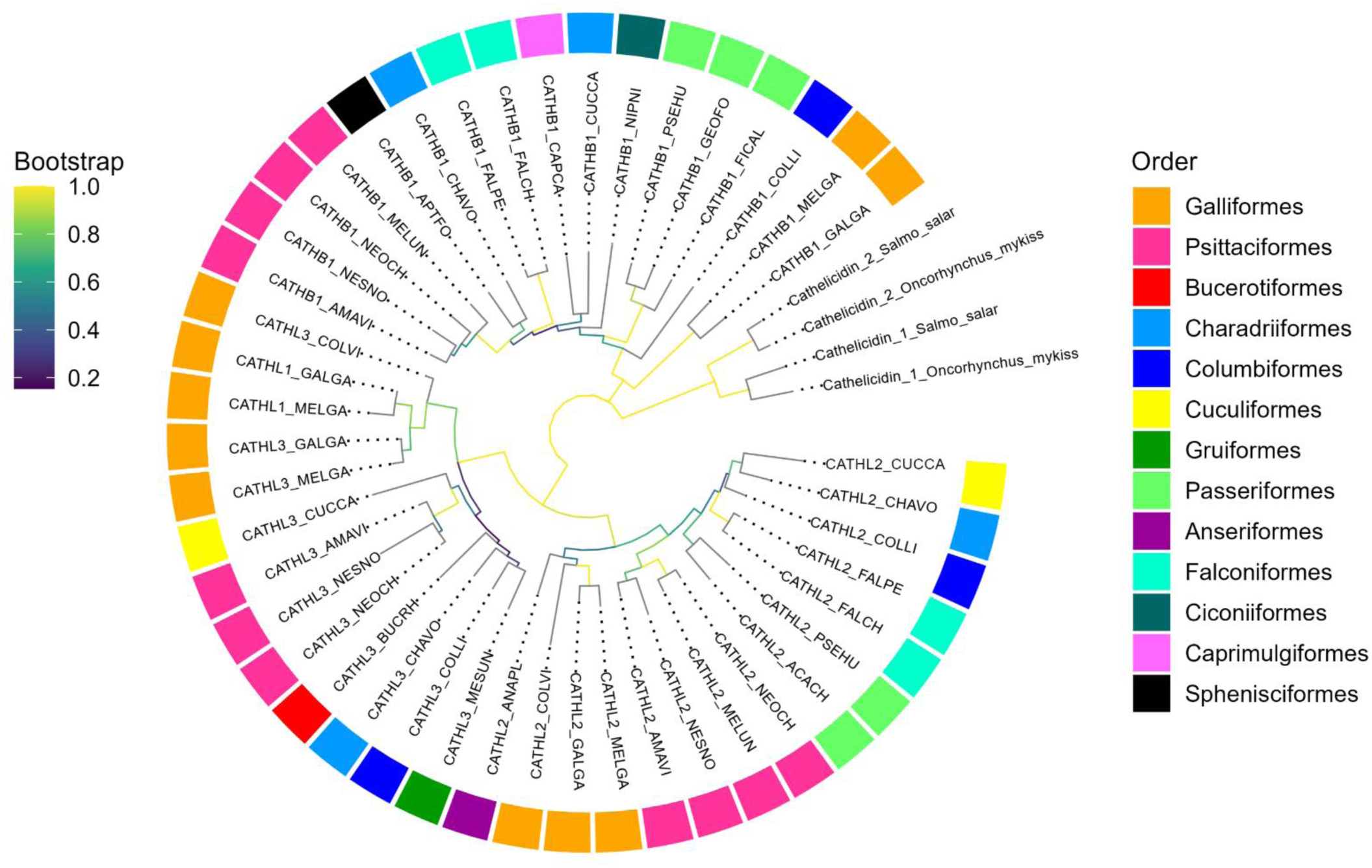
Neighbour-joining phylogenetic tree of full-length avian cathelicidins. The tree is rooted with fish cathelicidins. Branches are coloured according to bootstrap support. Tips are coloured according to order. Cathelicidins are labelled according to the nomenclature of (Cheng et al., 2015). For example, orange-bellied parrot cathelicidins are labelled CATHB1_NEOCH (*<u>Neo</u>phema <u>ch</u>rysogaster*).

**Figure S5:**
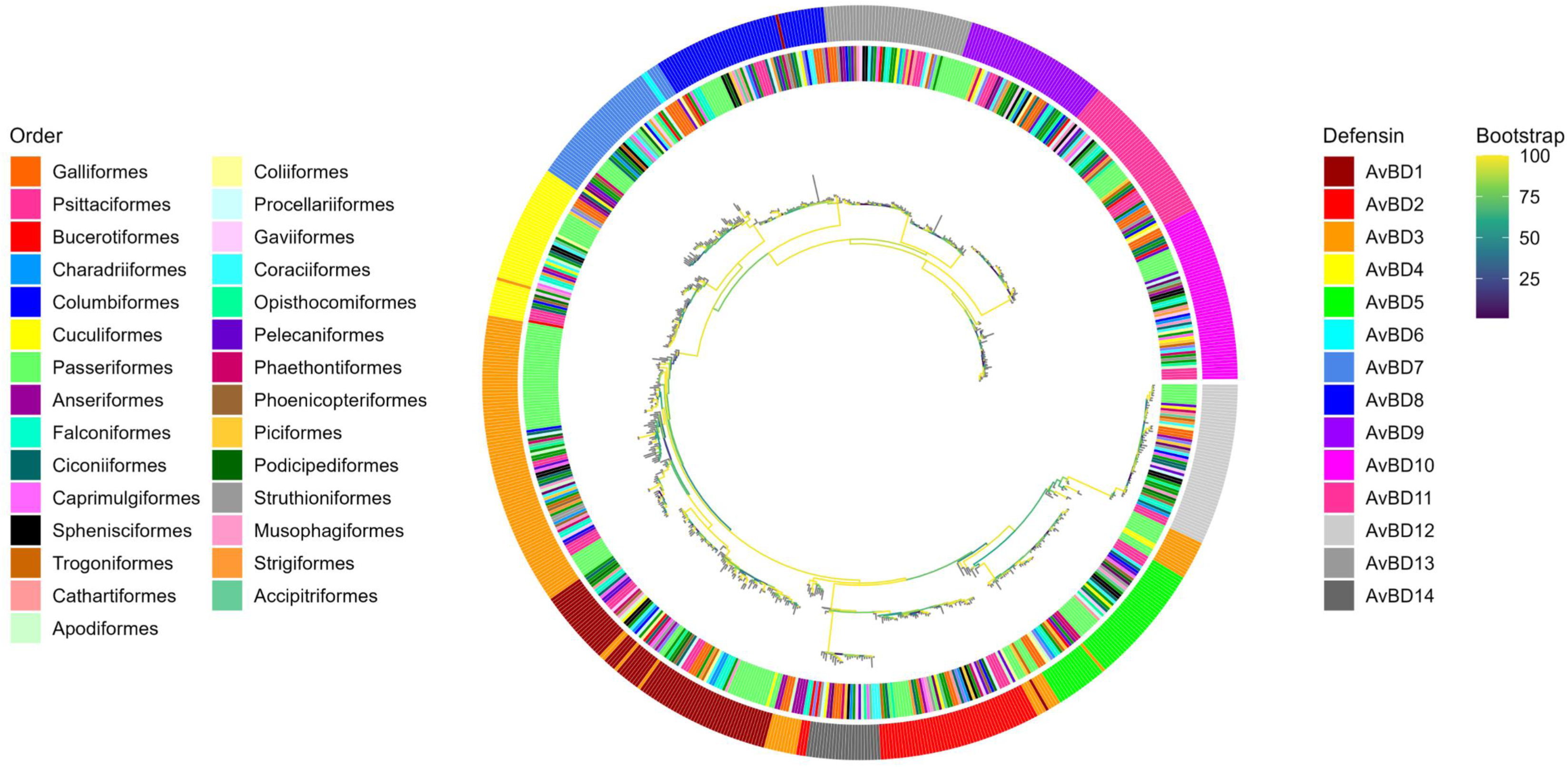
Maximum likelihood phylogenetic tree of full-length avian β-defensin amino acid sequences. The inner circle is coloured according to order. The outer circle is coloured according to β-defensin gene to show orthologous groups. Note, orange-bellied parrots sit within the Psittaciformes order.

**Figure S6:**
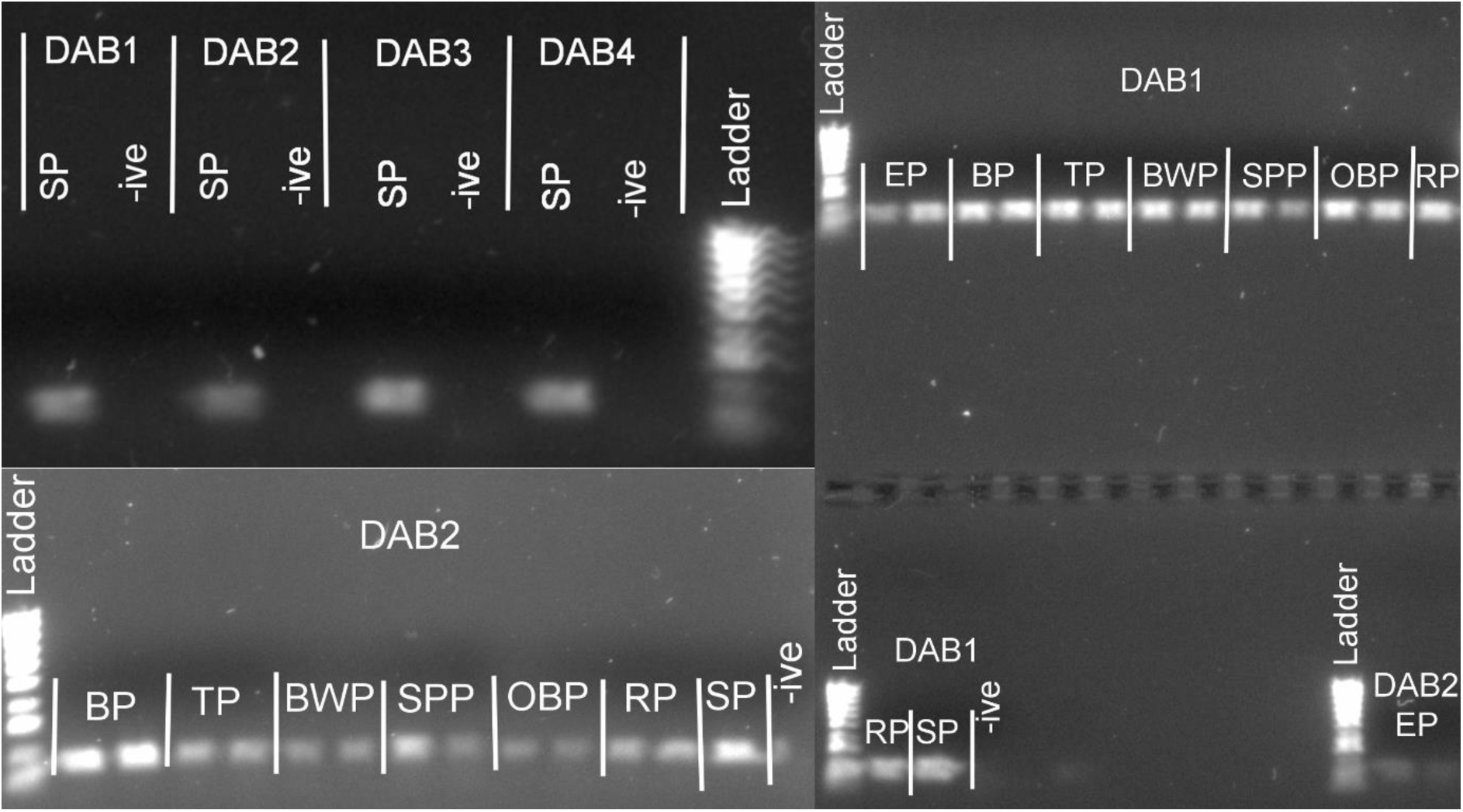
Gel image of MHC PCR amplification on a 1% agarose TBE gel stained with SYBR Safe DNA (Life Technologies) alongside Hyperladder IV (Bioline). Gel images show PCR amplification of MHC class II gene DAB in swift parrot (SP), elegant parrot (EP), Bourke’s parrot (BP), turquoise parrot (TP), blue-wing parrot (BWP), splendid parrot (SPP), orange bellied parrot (OBP) and rock parrot (RP) with four different sets of primers (DAB1, DAB2, DAB3, DAB4; see Table 2 for primers)

**Figure S7:**
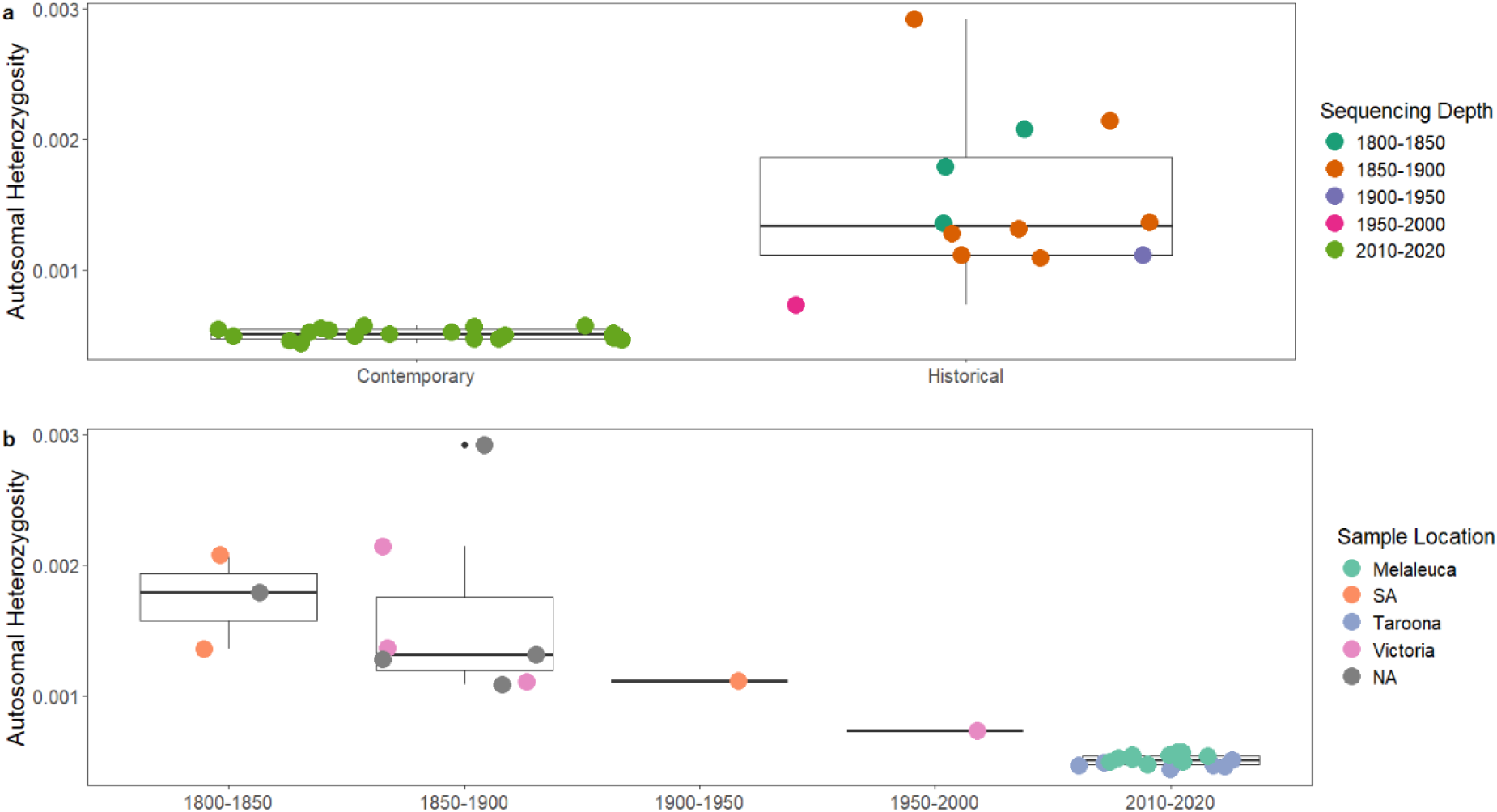
Autosomal heterozygosity as calculated by obtaining the site frequency spectrum using winSFS (v0.7.0) after all samples were downsampled to a 5x sequencing depth. Boxes represent the interquartile range (IQR) with whiskers extending to 1.5*IQR and bold line is the mean for each group.

**Figure S8:**
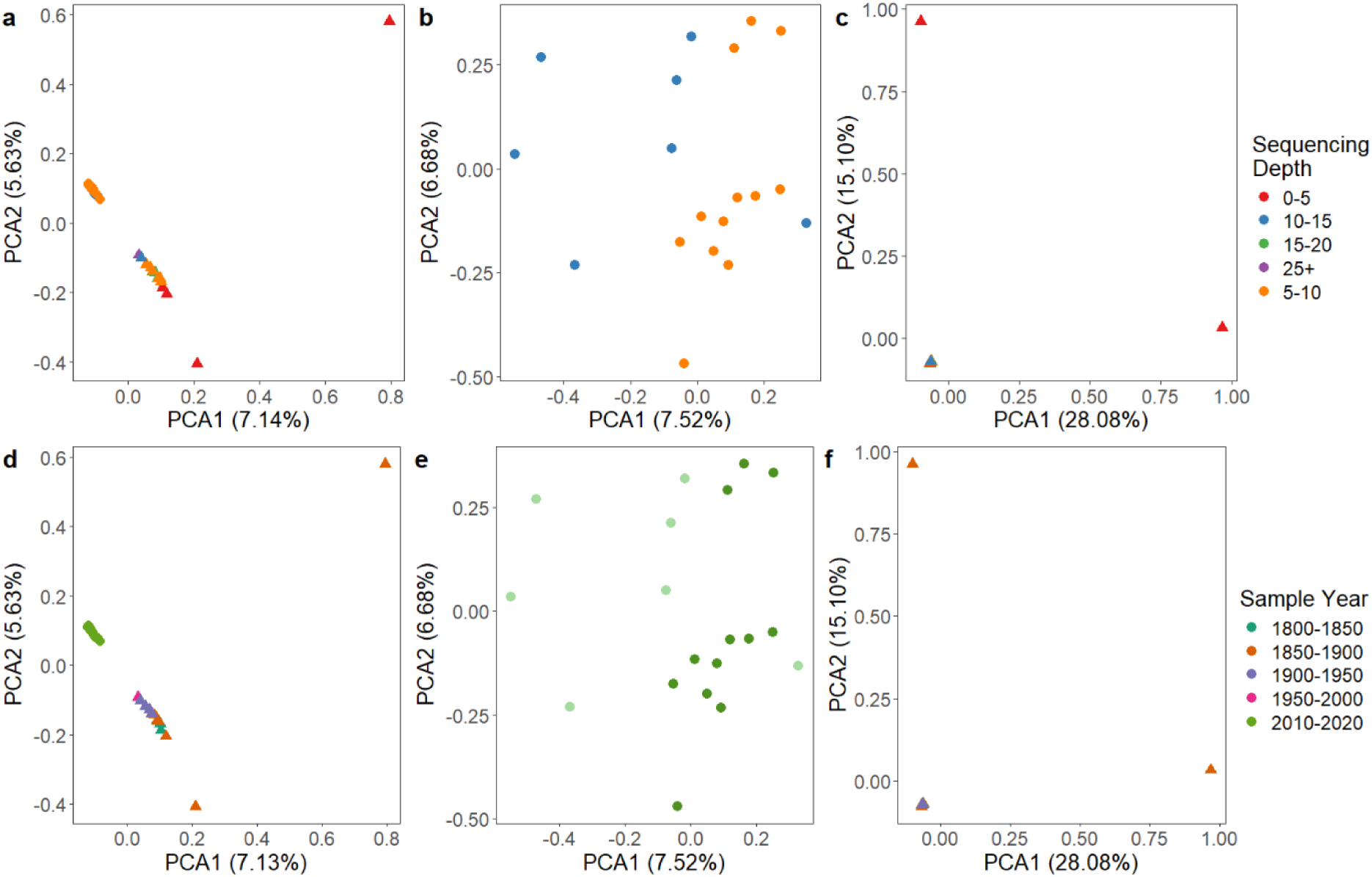
PCA plots of all (a, d), modern (b, e) and historical (c, f) samples calculated using angsd to output a covariance matrix for each group. Samples are coloured by sequencing depth (a, b, c) and sampling year (d, e, f). Modern samples are depicted with circles and historical samples by triangles. Note the two outliers in the historical samples (top left, bottom right) are causing all other historical samples to plot on top of one another.

**Figure S9:**
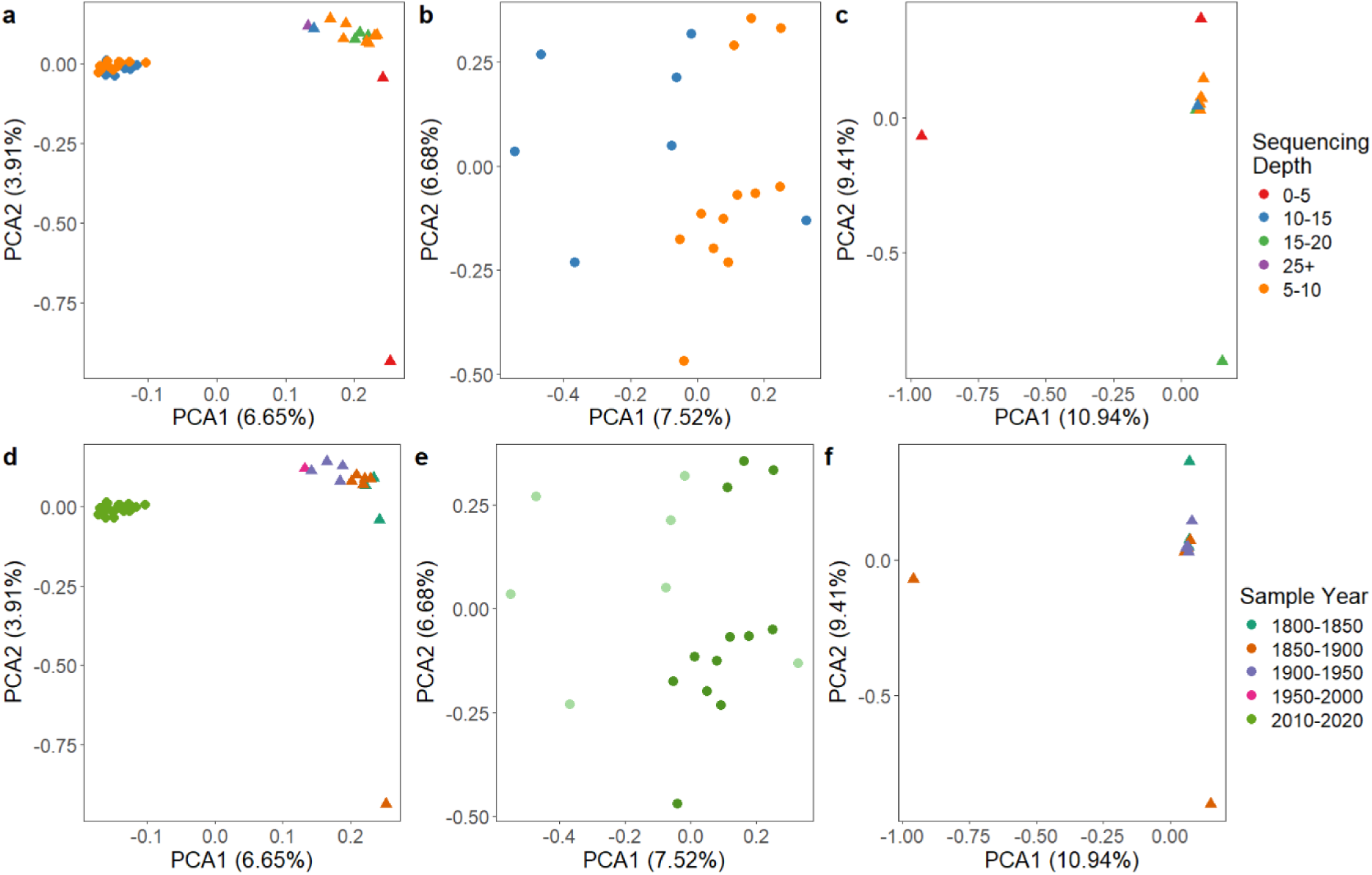
PCA plots of all (a, d), modern (b, e) and historical (c, f) samples (with the two outliers removed) calculated using angsd to output a covariance matrix for each group. Samples are coloured by sequencing depth (a, b, c) and sampling year (d, e, f). Modern samples are depicted with circles and historical samples by triangles.

**Figure S10:**
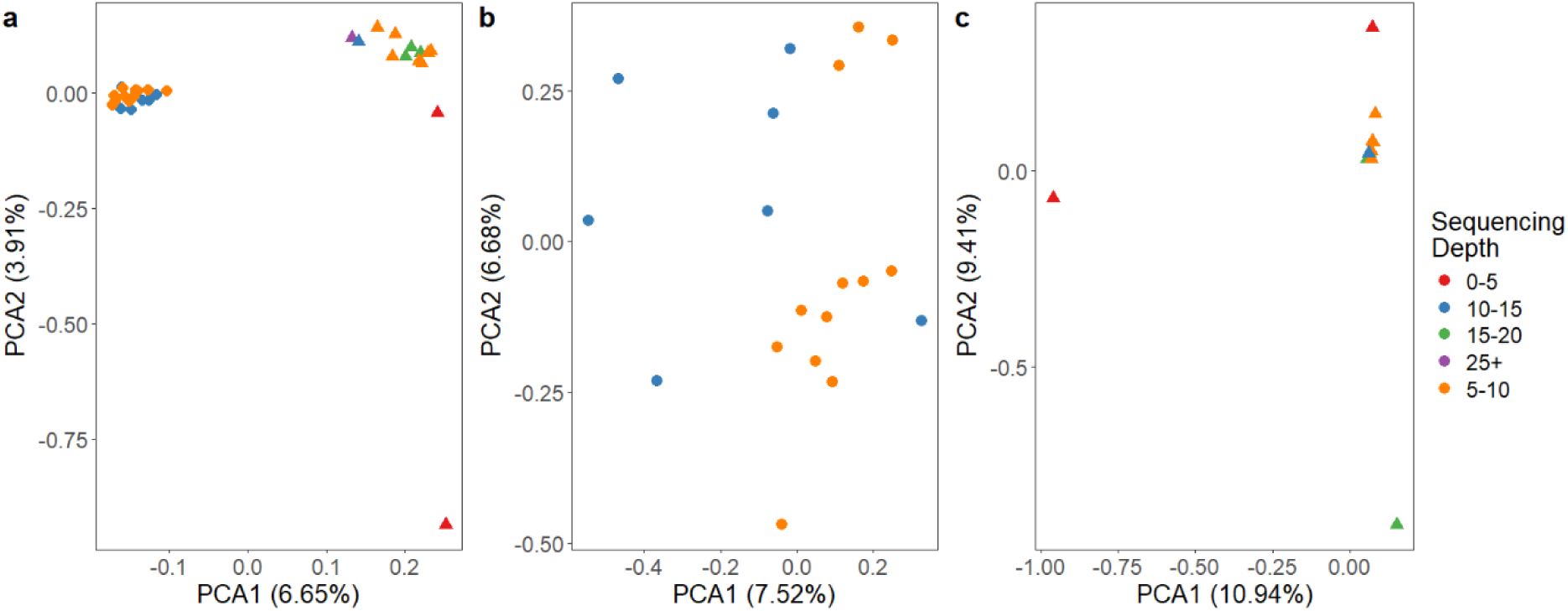
PCA plots of all (a), modern (b) and historical (c) samples calculated using angsd to output a covariance matrix for each group. Samples are coloured sampling year. Modern samples are depicted with circles and historical samples by triangles.

**Figure S11:**
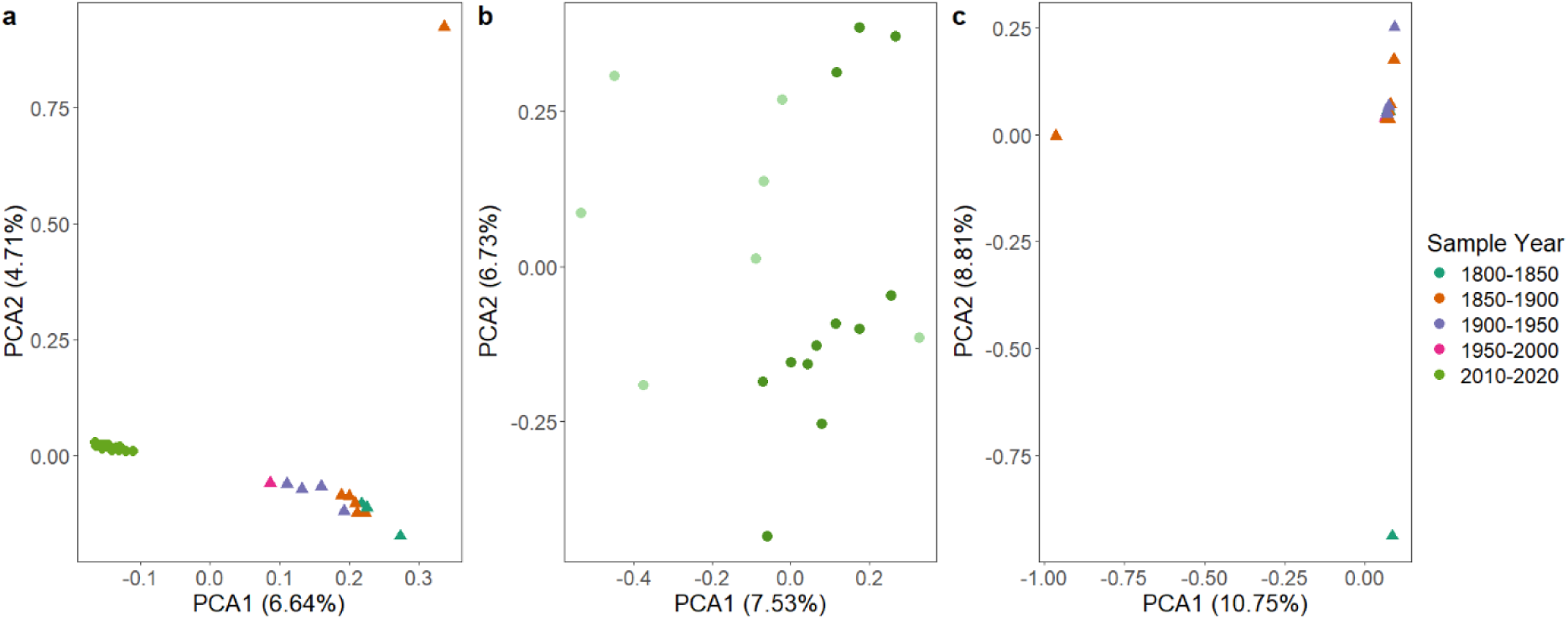
PCA plots of all (a), modern (b) and historical (c) samples calculated using angsd to output a covariance matrix for each group. Samples are coloured by sampling year. Modern samples are depicted with circles and historical samples by triangles.

**Figure S12:**
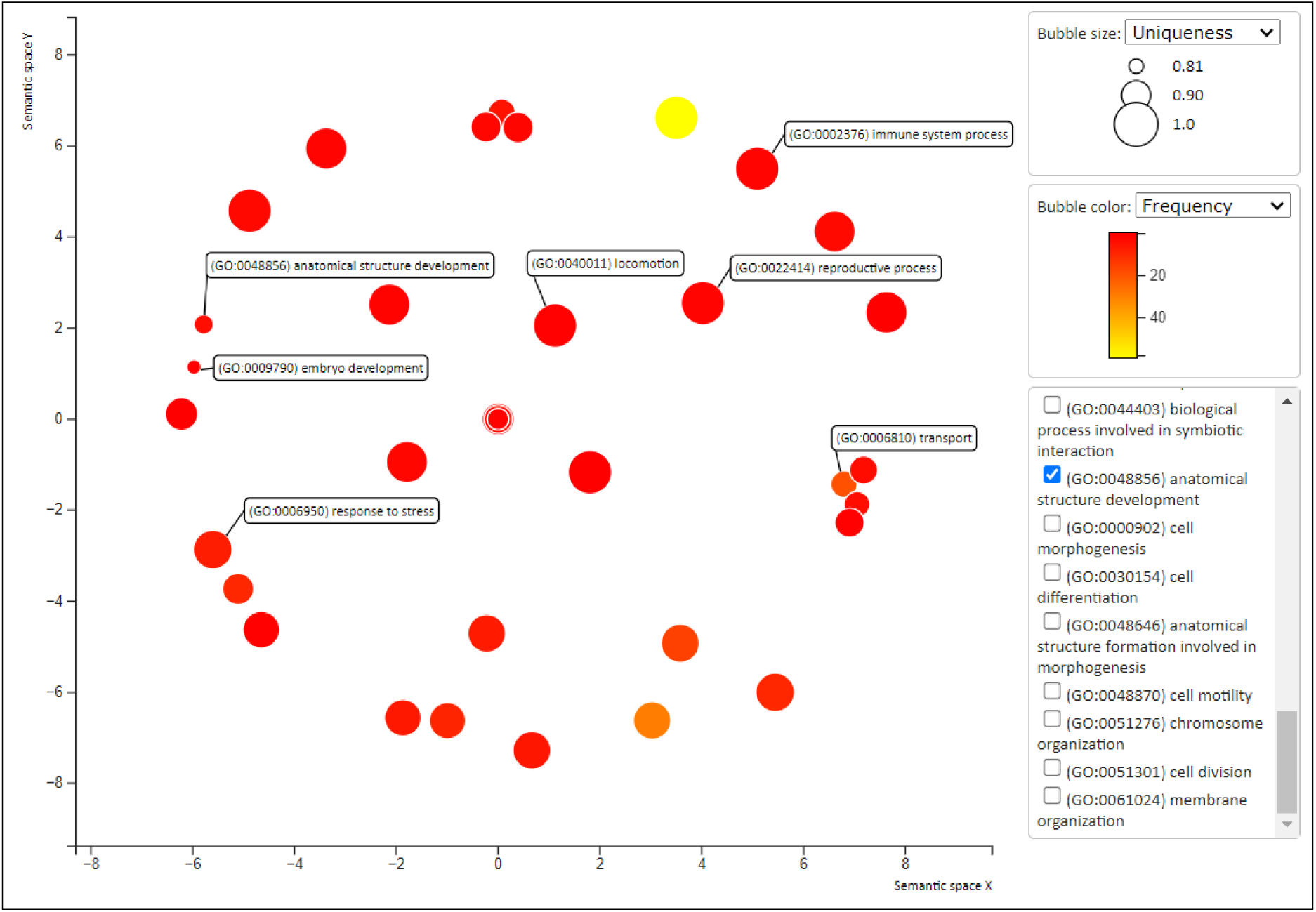
Clustering of GO terms by biological processes in semantic space by Revigo

**Figure S13:**
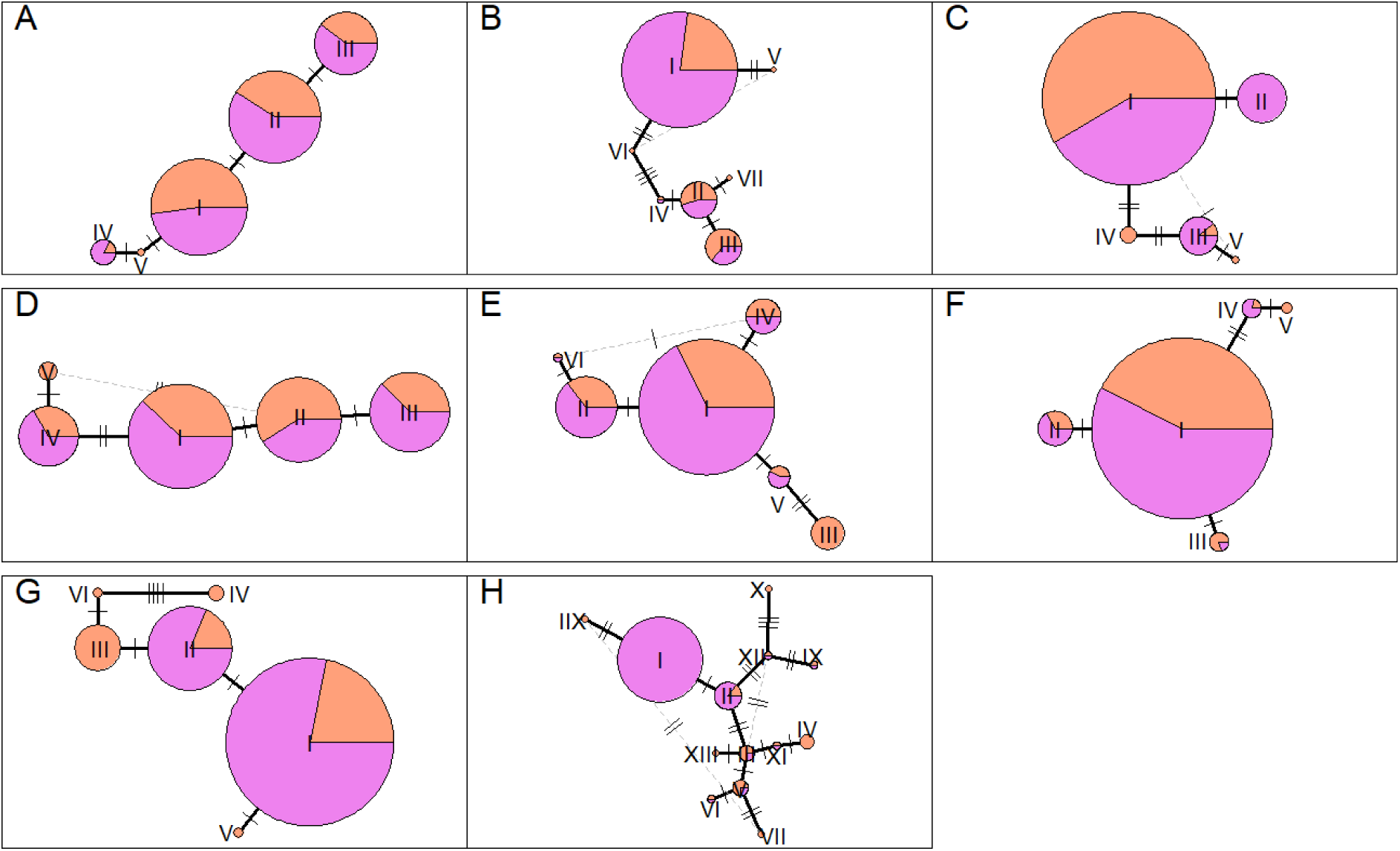
Allele network plots of TLR1A (A), TLR1B (B), TLR2A (C), TLR2B (D), TLR3 (E), TLR4 (F), TLR7 (G) and TLR15 (H) generated using pegas (v1.3) in R. Portions of each pie are represented by the proportion of contemporary (magenta) or historical (orange) samples with that allele.

**Table S1:**
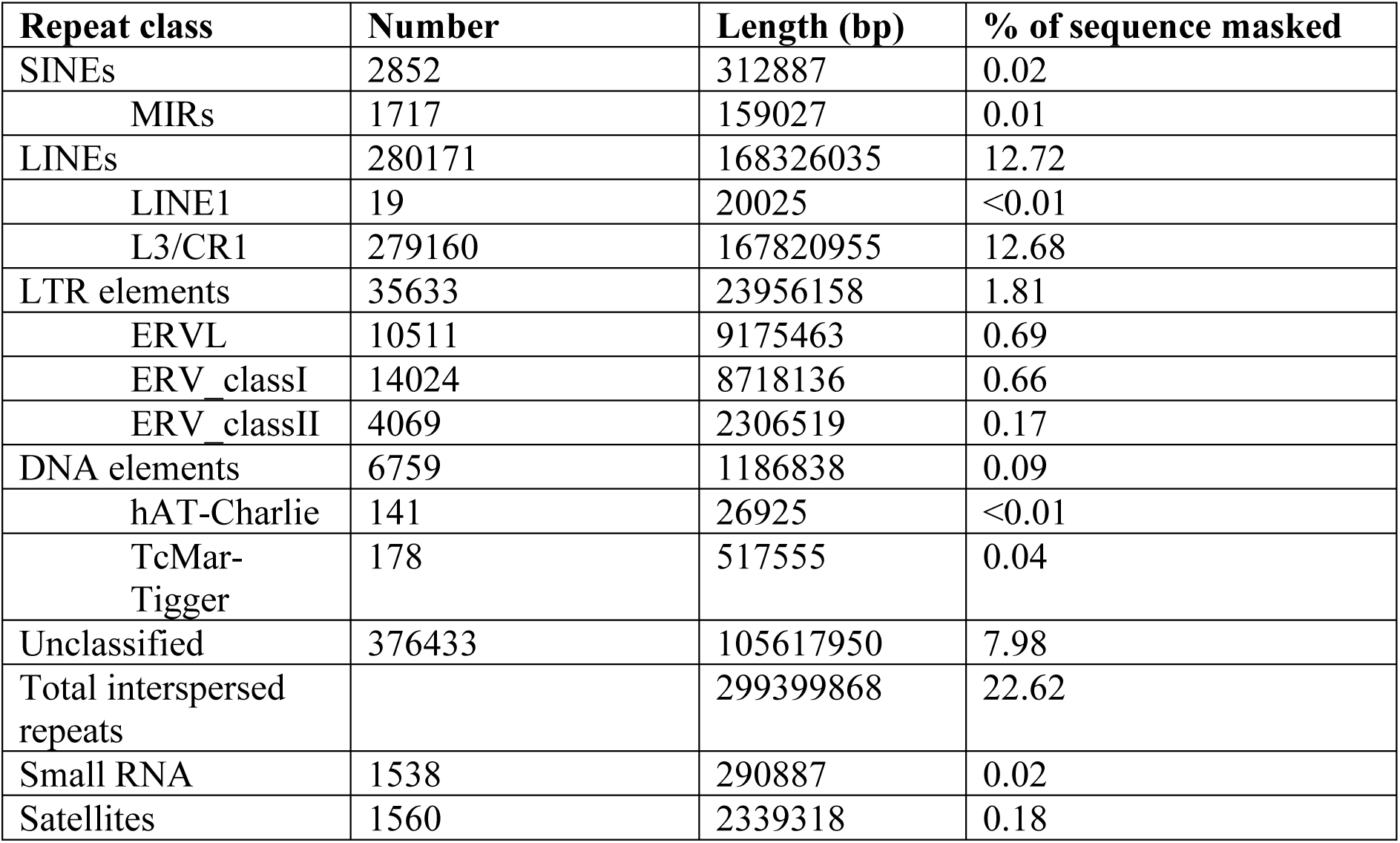
Repeats identified and masked in the genome assembly.

| **Repeat class** | **Number** | **Length (bp)** | **% of sequence masked** |
| --- | --- | --- | --- |
| SINEs | 2852 | 312887 | 0.02 |
| MIRs | 1717 | 159027 | 0.01 |
| LINEs | 280171 | 168326035 | 12.72 |
| LINE1 | 19 | 20025 | <0.01 |
| L3/CR1 | 279160 | 167820955 | 12.68 |
| LTR elements | 35633 | 23956158 | 1.81 |
| ERVL | 10511 | 9175463 | 0.69 |
| ERV_classI | 14024 | 8718136 | 0.66 |
| ERV_classII | 4069 | 2306519 | 0.17 |
| DNA elements | 6759 | 1186838 | 0.09 |
| hAT-Charlie | 141 | 26925 | <0.01 |
| TcMar-Tigger | 178 | 517555 | 0.04 |
| Unclassified | 376433 | 105617950 | 7.98 |
| Total interspersed repeats |  | 299399868 | 22.62 |
| Small RNA | 1538 | 290887 | 0.02 |
| Satellites | 1560 | 2339318 | 0.18 |

**Table S2:** QC and alignment of tissue transcriptomes. Percentage duplicated sequences and number of sequences (millions) in paired transcriptome reads post-trimming (averaged across the forward and reverse reads of the two lanes) and alignment rate to the repeat-masked genome.

| Bird | Tissue | Concentration (ng/ul) | RIN | Average % duplicated sequences (SD) | Total number of sequences (millions) | Alignment rate (%) |
| --- | --- | --- | --- | --- | --- | --- |
| 2035 | liver | 73.77 | 9.1 | 75.03 (0.005) | 98.6 | 91.15 |
| 2035 | spleen | 65.89 | 9.2 | 76.15 (0.006) | 124.6 | 91.11 |
| 2035 | kidney | 54.59 | 8.9 | 77.58 (0.004) | 111.0 | 90.91 |
| 2036 | pectoral muscle | 89.01 | 8.3 | 82.78 (0.007) | 116.4 | 88.34 |
| 1699 | heart | 75.44 | 8.9 | 74.90 (0.005) | 97.0 | 91.10 |
| 1699 | testis | 443.20 | 7.4 | 54.13 (0.009) | 110.6 | 84.60 |
| 1699 | tongue | 96.05 | 7.6 | 75.20 (0.006) | 101.4 | 88.86 |
| 1699 | pancreas | 129 | 8.0 | 73.45 (0.004) | 110.8 | 91.40 |

**Table S3:** BUSCO v5.5.0 scores of the global transcriptome assembly and genome annotation using aves_odb10 lineage dataset (n = 8338 BUSCOs).

| Assembly | Mode | Complete | Complete – single copy | Complete – duplicate | Fragmented | Missing |
| --- | --- | --- | --- | --- | --- | --- |
| Global transcriptome | Transcriptome | 91.5% | 12.8% | 78.7% | 2.2% | 6.3% |
| Annotation | Protein | 83.5% | 82.9% | 0.6% | 7.0% | 9.5% |

**Table S4:** Genomic coordinates of manually annotated immune genes..

| Scaffold | Feature | Start | End | Name | Strand |
| --- | --- | --- | --- | --- | --- |
| scaffold_1 | Gene | 24356825 | 24359282 | TLR1A | - |
| scaffold_1 | Exon | 24356825 | 24359282 | TLR1A_exon1 | - |
| scaffold_1 | Gene | 24366066 | 24368016 | TLR1B | - |
| scaffold_1 | Exon | 24366066 | 24368016 | TLR1B_exon1 | - |
| scaffold_1 | Gene | 34874661 | 34881447 | TLR3 | - |
| scaffold_1 | Exon | 34874661 | 34874890 | TLR3_exon4 | - |
| scaffold_1 | Exon | 34875562 | 34877382 | TLR3_exon3 | - |
| scaffold_1 | Exon | 34879401 | 34879593 | TLR3_exon2 | - |
| scaffold_1 | Exon | 34881006 | 34881447 | TLR3_exon1 | - |
| scaffold_1 | Gene | 47163399 | 47165751 | TLR2A | - |
| scaffold_1 | Exon | 47163399 | 47165751 | TLR2A_exon1 | - |
| scaffold_1 | Gene | 47171372 | 47179834 | TLR2B | - |
| scaffold_1 | Exon | 47171372 | 47173755 | TLR2B_exon3 | - |
| scaffold_1 | Exon | 47175508 | 47175585 | TLR2B_exon2 | - |
| scaffold_1 | Exon | 47179786 | 47179834 | TLR2B_exon1 | - |
| scaffold_16 | Gene | 4288411 | 4293031 | TLR4 | + |
| scaffold_16 | Exon | 4288411 | 4288588 | TLR4_exon1 | + |
| scaffold_16 | Exon | 4289402 | 4289569 | TLR4_exon2 | + |
| scaffold_16 | Exon | 4290771 | 4293031 | TLR4_exon3 | + |
| scaffold_3 | Gene | 76901499 | 76908664 | TLR7 | + |
| scaffold_3 | Exon | 76901499 | 76901502 | TLR7_exon1 | + |
| scaffold_3 | Exon | 76905523 | 76908664 | TLR7_exon2 | + |
| scaffold_4 | Gene | 1643032 | 1645654 | TLR15 | + |
| scaffold_4 | Exon | 1643032 | 1645654 | TLR15_exon1 | + |
| scaffold_4 | Gene | 108851307 | 108853896 | TLR5 | - |
| scaffold_4 | Exon | 108851307 | 108853896 | TLR5_exon1 | - |
| scaffold_174 | Gene | 198320 | 201139 | UA | + |
| scaffold_174 | Exon | 198320 | 198443 | UA_exon1 | + |
| scaffold_174 | Exon | 198910 | 199174 | UA_exon2 | + |
| scaffold_174 | Exon | 199294 | 199570 | UA_exon3 | + |
| scaffold_174 | Exon | 200197 | 200473 | UA_exon4 | + |
| scaffold_174 | Exon | 200553 | 200655 | UA_exon5 | + |
| scaffold_174 | Exon | 200756 | 200792 | UA_exon6 | + |
| scaffold_174 | Exon | 200933 | 200966 | UA_exon7 | + |
| scaffold_174 | Exon | 201125 | 201139 | UA_exon8 | + |
| scaffold_2 | Gene | 100078669 | 100079774 | CATHB1 | + |
| scaffold_2 | Exon | 100078669 | 100078916 | CATHB1_exon1 | + |
| scaffold_2 | Exon | 100079140 | 100079246 | CATHB1_exon2 | + |
| scaffold_2 | Exon | 100079353 | 100079437 | CATHB1_exon3 | + |
| scaffold_2 | Exon | 100079570 | 100079774 | CATHB1_exon4 | + |
| scaffold_2 | Gene | 100080112 | 100080975 | CATHL3 | + |
| scaffold_2 | Exon | 100080112 | 100080285 | CATHL3_exon1 | + |
| scaffold_2 | Exon | 100080496 | 100080615 | CATHL3_exon2 | + |
| scaffold_2 | Exon | 100080720 | 100080800 | CATHL3_exon3 | + |
| scaffold_2 | Exon | 100080877 | 100080975 | CATHL3_exon4 | + |
| scaffold_2 | Gene | 100083118 | 100082023 | CATHL2 | - |
| scaffold_2 | Exon | 100083118 | 100082912 | CATHL2_exon1 | - |
| scaffold_2 | Exon | 100082832 | 100082722 | CATHL2_exon2 | - |
| scaffold_2 | Exon | 100082621 | 100082537 | CATHL2_exon3 | - |
| scaffold_2 | Exon | 100082143 | 100082023 | CATHL2_exon4 | - |
| scaffold_4 | Gene | 17394461 | 17395082 | AvBD1 | - |
| scaffold_4 | Exon | 17394461 | 17394691 | AvBD1_exon1 | - |
| scaffold_4 | Exon | 17394924 | 17395082 | AvBD1_exon2 | - |
| scaffold_4 | Gene | 17383047 | 17383828 | AvBD2 | + |
| scaffold_4 | Exon | 17383047 | 17383206 | AvBD2_exon1 | + |
| scaffold_4 | Exon | 17383604 | 17383828 | AvBD2_exon2 | + |
| scaffold_4 | Gene | 17399065 | 17401202 | AvBD3_1 | - |
| scaffold_4 | Exon | 17399065 | 17399281 | AvBD3_exon1 | - |
| scaffold_4 | Exon | 17401042 | 17401202 | AvBD3_exon2 | - |
| scaffold_4 | Gene | 17404250 | 17405332 | AvBD3_2 | - |
| scaffold_4 | Exon | 17404250 | 17404548 | AvBD3_2_exon1 | - |
| scaffold_4 | Exon | 17405172 | 17405332 | AvBD3_2_exon2 | - |
| scaffold_4 | Gene | 17408400 | 17409462 | AvBD3_3 | - |
| scaffold_4 | Exon | 17408400 | 17408678 | AvBD3_3_exon1 | - |
| scaffold_4 | Exon | 17409302 | 17409462 | AvBD3_3_exon2 | - |
| scaffold_4 | Gene | 17418212 | 17420540 | AvBD4 | - |
| scaffold_4 | Exon | 17418212 | 17418442 | AvBD4_exon1 | - |
| scaffold_4 | Exon | 17420382 | 17420540 | AvBD4_exon2 | - |
| scaffold_4 | Gene | 17413119 | 17413966 | AvBD5 | - |
| scaffold_4 | Exon | 17413119 | 17413349 | AvBD5_exon1 | - |
| scaffold_4 | Exon | 17413803 | 17413966 | AvBD5_exon2 | - |
| scaffold_4 | Gene | 17372537 | 17373138 | AvBD8 | - |
| scaffold_4 | Exon | 17372537 | 17372771 | AvBD8_exon1 | - |
| scaffold_4 | Exon | 17372979 | 17373138 | AvBD8_exon2 | - |
| scaffold_4 | Gene | 17365616 | 17368619 | AvBD9 | + |
| scaffold_4 | Exon | 17365616 | 17365776 | AvBD9_exon1 | + |
| scaffold_4 | Exon | 17368386 | 17368619 | AvBD9_exon2 | + |
| scaffold_4 | Gene | 17357081 | 17357615 | AvBD10 | + |
| scaffold_4 | Exon | 17357081 | 17357240 | AvBD10_exon1 | + |
| scaffold_4 | Exon | 17357379 | 17357615 | AvBD10_exon2 | + |
| scaffold_4 | Gene | 17334583 | 17336764 | AvBD11 | - |
| scaffold_4 | Exon | 17334583 | 17334458 | AvBD11_exon1 | - |
| scaffold_4 | Exon | 17335556 | 17335787 | AvBD11_exon2 | - |
| scaffold_4 | Exon | 17336615 | 17336764 | AvBD11_exon3 | - |
| scaffold_4 | Gene | 17327408 | 17328921 | AvBD12 | - |
| scaffold_4 | Exon | 17327408 | 17327599 | AvBD12_exon1 | - |
| scaffold_4 | Exon | 17328971 | 17328921 | AvBD12_exon2 | - |
| scaffold_4 | Gene | 17319021 | 17320610 | AvBD13 | - |
| scaffold_4 | Exon | 17319021 | 17319227 | AvBD13_exon1 | - |
| scaffold_4 | Exon | 17320450 | 17320610 | AvBD13_exon2 | - |

**Table S5:** Sample and sequencing metadata for the resequenced genomes.

| Sample | Museum Code | Institution | Year | Autosomal heterozygosity | No. reads | Mapped reads | Endogenous % | Coverage | Depth | % unambiguous realign |
| --- | --- | --- | --- | --- | --- | --- | --- | --- | --- | --- |
| O-76339-001 |  | Australian Museum | 2016 | 0.00058914 | 67,774,977 | 67,370,930 | 99.4 | 86.50 | 13.42 | 74.31 |
| O-76349-001 |  | Australian Museum | 2016 | 0.00053986 | 70,854,359 | 70,636,998 | 99.69 | 86.39 | 14.58 | 86.76 |
| O-76350-001 |  | Australian Museum | 2016 | 0.00055870 | 67,450,912 | 67,267,333 | 99.73 | 85.84 | 13.03 | 90.94 |
| O-76521-001 |  | Australian Museum | 2016 | 0.00055385 | 67,577,976 | 67,161,657 | 99.38 | 85.67 | 13.37 | 77.54 |
| O-76523-001 |  | Australian Museum | 2016 | 0.00057339 | 67,730,944 | 67,526,420 | 99.7 | 86.22 | 13.99 | 88.69 |
| O-76526-001 |  | Australian Museum | 2016 | 0.00056664 | 67,579,171 | 67,385,460 | 99.71 | 86.84 | 14.45 | 88.68 |
| O-76538-001 |  | Australian Museum | 2016 | 0.00060983 | 67,408,092 | 67,190,117 | 99.68 | 86.55 | 13.91 | 87.27 |
| O-81171-001 |  | Australian Museum | 2020 | 0.00059708 | 67,548,302 | 67,371,596 | 99.74 | 78.57 | 6.30 | 89.62 |
| O-81175-001 |  | Australian Museum | 2020 | 0.00054620 | 67,439,419 | 67,093,654 | 99.49 | 78.58 | 6.33 | 79.30 |
| O-81179-001 |  | Australian Museum | 2020 | 0.00054062 | 69,198,681 | 69,017,732 | 99.74 | 80.28 | 7.87 | 89.06 |
| O-81182-001 |  | Australian Museum | 2020 | 0.00056421 | 67,136,696 | 66,969,341 | 99.75 | 75.44 | 5.44 | 91.30 |
| O-81186-001 |  | Australian Museum | 2020 | 0.00060028 | 65,865,224 | 65,546,200 | 99.52 | 74.37 | 5.27 | 82.73 |
| O-81188-001 |  | Australian Museum | 2020 | 0.00055175 | 67,559,256 | 67,178,228 | 99.44 | 79.42 | 6.87 | 80.87 |
| O-81192-001 |  | Australian Museum | 2020 | 0.00056527 | 67,710,387 | 67,381,578 | 99.51 | 75.49 | 5.70 | 78.48 |
| O-81193-001 |  | Australian Museum | 2020 | 0.00060828 | 67,387,149 | 67,225,623 | 99.76 | 75.76 | 5.32 | 89.76 |
| O-81196-001 |  | Australian Museum | 2020 | 0.00063218 | 67,358,970 | 66,996,167 | 99.46 | 78.53 | 6.74 | 81.60 |
| O-81198-001 |  | Australian Museum | 2020 | 0.00057471 | 57,844,720 | 57,709,356 | 99.77 | 74.39 | 5.20 | 91.84 |
| O-81201-001 |  | Australian Museum | 2020 | 0.00059226 | 67,104,272 | 66,946,028 | 99.76 | 75.51 | 5.44 | 91.10 |
| O-81203-001 |  | Australian Museum | 2020 | 0.00055296 | 67,303,297 | 67,148,173 | 99.77 | 76.03 | 5.70 | 92.20 |
| OBPI_501 | B17317 | Museums Victoria | 1986 | 0.00071108 | 401,292,042 | 339,034,474 | 84.49 | 98.21 | 25.32 | 73.83 |
| OBPI_502 | R8767 | Museums Victoria | 1881 | 0.00087726 | 335,811,013 | 188,636,808 | 56.17 | 97.98 | 15.28 | 66.37 |
| OBPI_503 | R8772 | Museums Victoria | 1884 | 0.00206197 | 190,838,260 | 125,800,628 | 65.92 | 97.13 | 17.18 | 71.23 |
| OBPI_504 | R8773 | Museums Victoria | 1884 | 0.00082324 | 551,691,763 | 319,811,217 | 57.97 | 98.28 | 17.83 | 75.54 |
| OBPI_505 | 1843.7.14.91 | South Australian Museum | 1843 | 0.00151453 | 130,192,767 | 54,498,918 | 41.86 | 71.99 | 4.07 | 21.15 |
| OBPI_506 | 1843.7.14.92 | South Australian Museum | 1843 | 0.00132283 | 221,495,144 | 99,556,628 | 44.95 | 87.08 | 6.44 | 51.58 |
| OBPI_507 | 1845.11.7.9 | South Australian Museum | 1845 | 0.00151321 | 293,602,567 | 125,045,402 | 42.59 | 93.63 | 8.93 | 23.06 |
| OBPI_508 | 1856.3.14.43 | South Australian Museum | 1856 | 0.00125715 | 143,662,086 | 81,030,754 | 56.4 | 74.73 | 4.28 | 42.69 |

| **Sample** | **Museum Code** | **Institution** | **Year** | **Autosomal heterozygosity** | **No. reads** | **Mapped reads** | **Endogenous %** | **Coverage** | **Depth** | **% unambiguous realign** |
| --- | --- | --- | --- | --- | --- | --- | --- | --- | --- | --- |
| OBPII_Re HoI_501 | 1881.5.1.4796 | South Australian Museum | 1881 | 0.00188992 | 230,976,155 | 106,933,216 | 46.3 | 91.74 | 6.64 | 32.21 |
| OBPII_Re HoI_502 | 1996-617 | Paris | 1887 | 0.00126975 | 210,448,396 | 128,787,082 | 61.2 | 89.84 | 6.22 | 61.22 |
| OBPII_Re HoI_503 | 2004-27 | Paris | 1829 | 0.00261626 | 188,306,288 | 84,431,824 | 44.84 | 42.74 | 2.48 | 40.44 |
| OBPII_Re HoI_504 | B2387 | South Australian Museum | 1918 | 0.00087820 | 217,126,835 | 139,868,322 | 64.42 | 95.12 | 7.75 | 63.28 |
| OBPII_Re HoI_505 | B3025 | South Australian Museum | 1863 | 0.00225285 | 134,019,630 | 58,956,678 | 43.99 | 71.63 | 2.92 | 32.57 |
| OBPII_Re HoI_506 | 0.35544.001 | South Australian Museum | 1937 | 0.00302827 | 159,771,733 | 85,678,249 | 53.63 | 90.18 | 5.59 | 24.72 |
| OBPII_Re HoI_507 | B11067 | South Australian Museum | 1927 | 0.00101881 | 259,935,350 | 169,230,871 | 65.1 | 96.1821 | 8.82 | 64.04 |
| OBPII_Re HoI_508 | B11068 | South Australian Museum | 1927 | 0.00080873 | 269,220,549 | 184,872,265 | 68.67 | 96.65 | 10.06 | 58.64 |

**Table S6:**
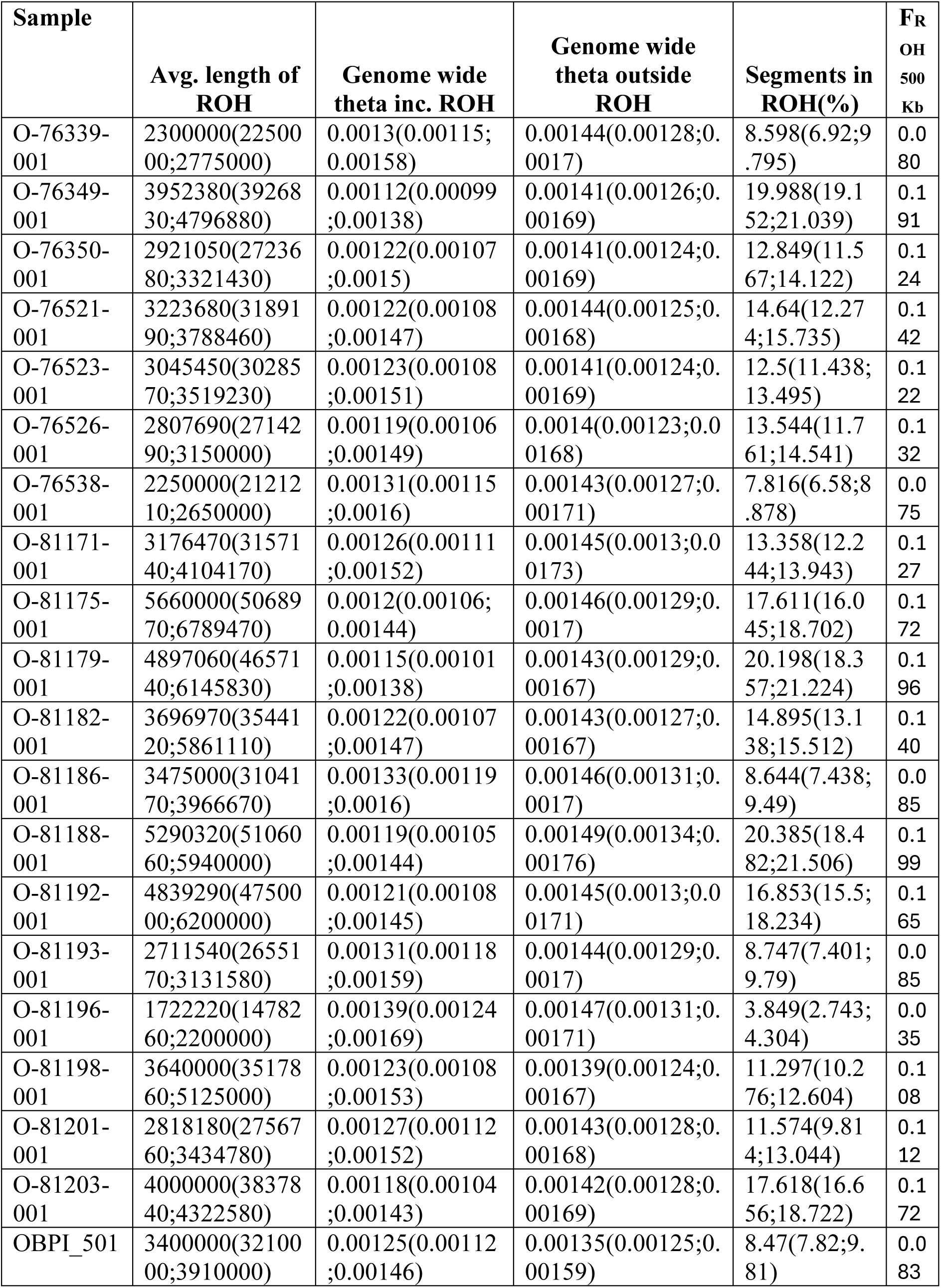
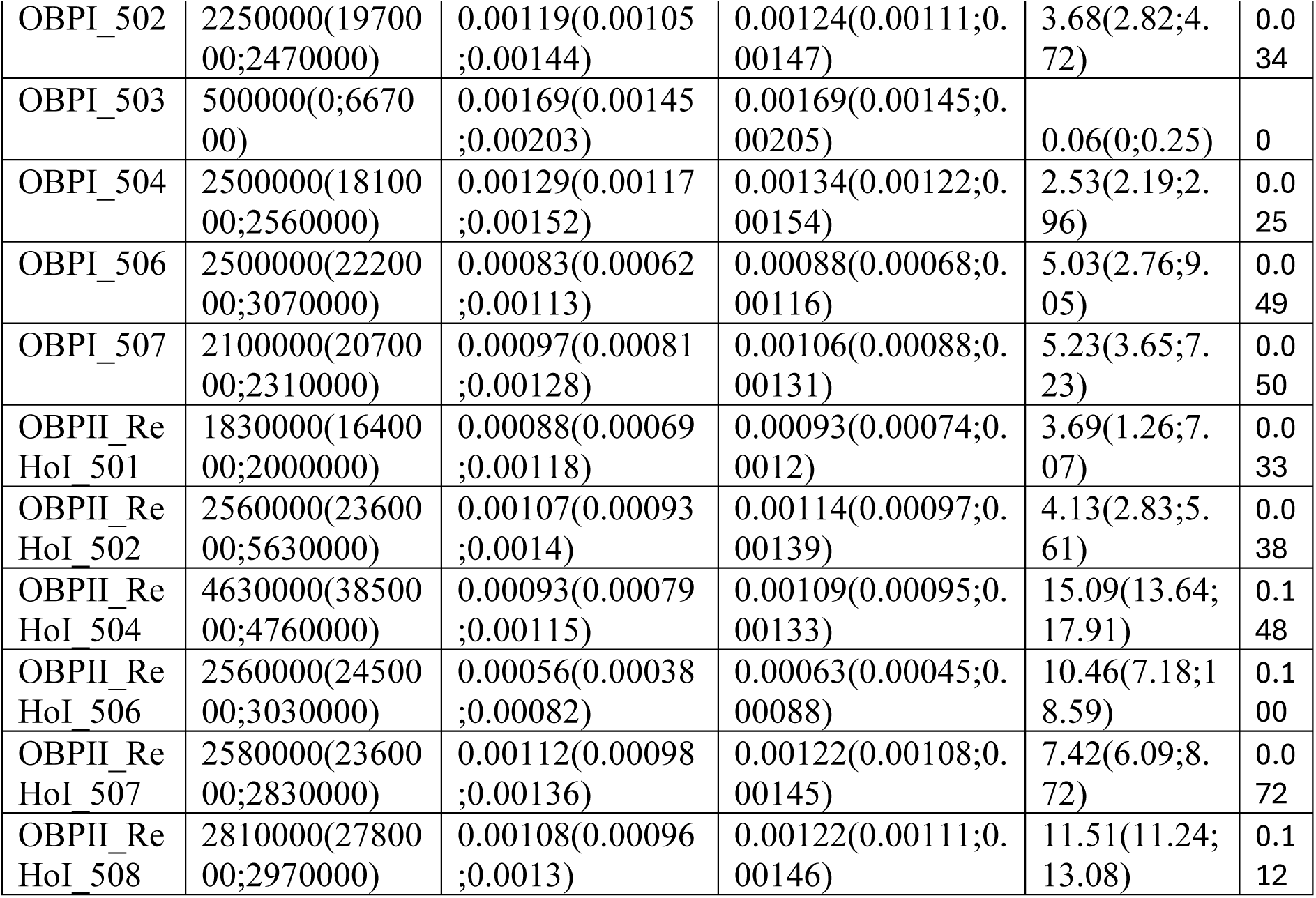
Runs of homozygosity (ROH) in modern and historic genomes. Percentage of autosomal genome in ROH and estimates of genome-wide Watterson’s theta with or without regions in ROH. Values calculated using a minimum ROH length of 500kb, rohmu of 5e-5 and TsTv ratio of 3.

**Table S7:** Gene ontology terms of 135 genes covered by ROHs in multiple individuals as determined by GoNet.

|  | GO Term ID | GO Term Definition | Genes |
| --- | --- | --- | --- |
| 1 | GO:0048856 | anatomical structure development | CHRD\|CLCN2\|DVL3\|EPHB3\|FETUA\|FUT9\|HRG\|HSPB7\|JIP3\|RAG1\|RAG2\|ROGDI\|TPO\|UFL1 |
| 2 | GO:0006464 | cellular protein modification process | ALG3\|CSN7B\|EPHB3\|FETUA\|FUT9\|HERC4\|JIP3\|KNG1\|PSMD2\|RAG1\|RAG2\|SENP5\|UFL1 |
| 3 | GO:0006950 | response to stress | ABCF3\|CSN7B\|FETUA\|HRG\|HSPB7\|JIP3\|KNG1\|PSMD2\|RAG2\|TPO\|UFL1 |
| 4 | GO:0006810 | transport | ANO2\|AP2M1\|CLCN2\|FETUA\|HRG\|JIP3\|KNG1\|MRP5\|NMUR1\|PSMD2 |
| 5 | GO:0030154 | cell differentiation | CHRD\|CLCN2\|DVL3\|EPHB3\|HERC4\|JIP3\|RAG1\|RAG2\|ROGDI\|UFL1 |
| 6 | GO:0002376 | immune system process | ABCF3\|AP2M1\|EPHB3\|FETUA\|HRG\|PSMD2\|RAG1\|RAG2\|ROGDI\|TPO |
| 7 | GO:0007165 | signal transduction | AP2M1\|CHRD\|DVL3\|ECEL1\|EPHB3\|GPR63\|JIP3\|KNG1\|NMUR1\|PSMD2 |
| 8 | GO:0016192 | vesicle-mediated transport | AP2M1\|FETUA\|HRG\|JIP3\|KNG1\|PSMD2 |
| 9 | GO:0009058 | biosynthetic process | ALG3\|FUT9\|MRP5\|PTMA\|TELO2\|TPO |
| 10 | GO:0051276 | chromosome organization | CSN7B\|PTMA\|RAG1\|RAG2\|TELO2 |
| 11 | GO:0042592 | homeostatic process | KNG1\|RAG1\|RAG2\|ROGDI\|TELO2 |
| 12 | GO:0034641 | cellular nitrogen compound metabolic process | CSN7B\|PTMA\|RAG1\|RAG2\|TELO2 |
| 13 | GO:0006259 | DNA metabolic process | CSN7B\|RAG1\|RAG2\|TELO2 |
| 14 | GO:0055085 | transmembrane transport | ANO2\|CLCN2\|MRP5\|PSMD2 |
| 15 | GO:0048646 | anatomical structure formation involved in morphogenesis | CHRD\|DVL3\|EPHB3\|HRG |
| 16 | GO:0009056 | catabolic process | AP2M1\|FUT9\|PSMD2\|TPO |
| 17 | GO:0022607 | cellular component assembly | AP2M1\|ARMC9\|CSN7B |
| 18 | GO:0007267 | cell-cell signaling | AP2M1\|DVL3\|PSMD2 |
| 19 | GO:0009790 | embryo development | CHRD\|DVL3\|TPO |
| 20 | GO:0040011 | locomotion | EPHB3\|HRG |
| 21 | GO:0044281 | small molecule metabolic process | FUT9\|MRP5 |

|  | **GO Term ID** | **GO Term Definition** | **Genes** |
| --- | --- | --- | --- |
| 22 | GO:0050877 | nervous system process | PDE6D\|RAG1 |
| 23 | GO:0015031 | protein transport | AP2M1\|JIP3 |
| 24 | GO:0065003 | protein-containing complex assembly | AP2M1\|CSN7B |
| 25 | GO:0044403 | symbiotic process | AP2M1\|CSN7B |
| 26 | GO:0008283 | cell population proliferation | RAG2 |
| 27 | GO:0030705 | cytoskeleton-dependent intracellular transport | JIP3 |
| 28 | GO:0048870 | cell motility | EPHB3 |
| 29 | GO:0051186 | cofactor metabolic process | TPO |
| 30 | GO:0051301 | cell division | SENP5 |
| 31 | GO:0061024 | membrane organization | AP2M1 |
| 32 | GO:0000003 | reproduction | HERC4 |
| 33 | GO:0007155 | cell adhesion | EPHB3 |
| 34 | GO:0006629 | lipid metabolic process | ALG3 |
| 35 | GO:0000902 | cell morphogenesis | EPHB3 |
| 36 | GO:0007049 | cell cycle | SENP5 |
| 37 | GO:0003013 | circulatory system process | KNG1 |
| 38 | GO:0005975 | carbohydrate metabolic process | FUT9 |

**Table S8:** Sites identified to be under selection using FEL on Datamonkey server.

| **Gene** | **Codon** | **alpha** | **beta** | **alpha=beta** | **LRT** | **▲ p-value** | **Total branch length** | **class** |
| --- | --- | --- | --- | --- | --- | --- | --- | --- |
| TLR1A | 120 | 299 | 0 | 64.46 | 2.735 | 0.0982 | 0.208 | Purifying |
| TLR1B | 54 | 294 | 0 | 102.1 | 4.2 | 0.0404 | 1.025 | Purifying |
| TLR1B | 223 | 217 | 0 | 76 | 3.979 | 0.0461 | 0.763 | Purifying |
| TLR1B | 192 | 215 | 0 | 39.16 | 2.82 | 0.0931 | 0.393 | Purifying |
| TLR3 | 117 | 692 | 0 | 85.39 | 3.808 | 0.051 | 0.315 | Purifying |
| TLR3 | 700 | 306 | 0 | 66.01 | 2.968 | 0.0849 | 0.244 | Purifying |
| TLR7 | 961 | 515 | 0 | 61.04 | 3.673 | 0.0553 | 0.375 | Purifying |
| TLR7 | 435 | 219 | 0 | 50.35 | 2.817 | 0.0933 | 0.309 | Purifying |
| TLR7 | 501 | 266 | 0 | 60.16 | 2.812 | 0.0936 | 0.37 | Purifying |

**Table S9:**
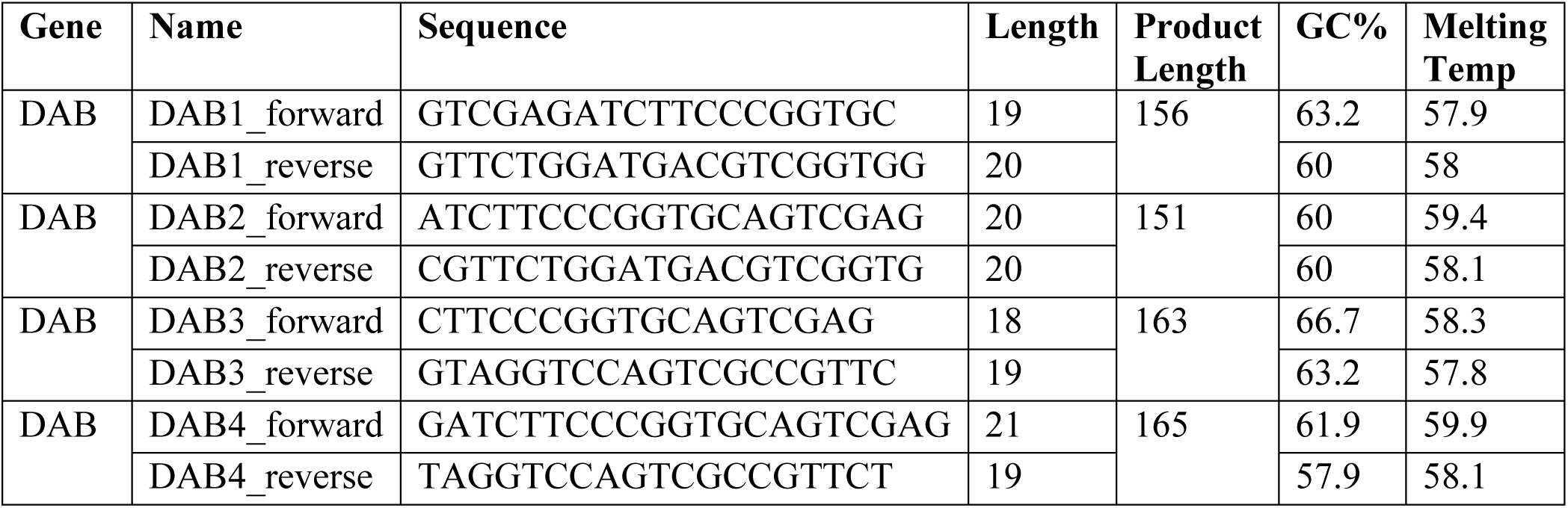
Sequences designed to amplify exon 2 of MHC genes in parrots.

**Table S10:**
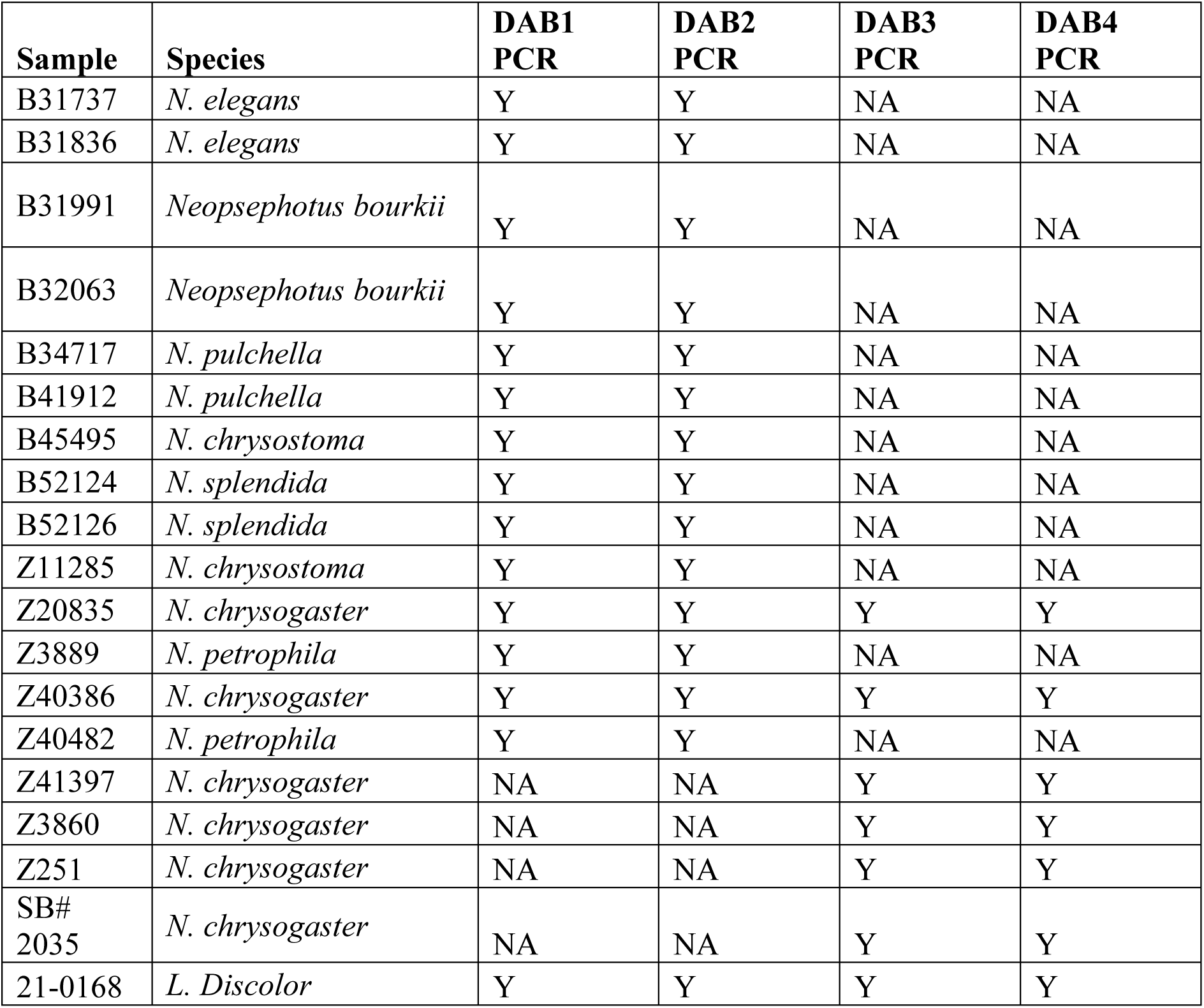
Sample ID of samples used for PCR analysis, Y indicates amplification of the MHC fragment occurred, NA indicates PCR was not performed on this sample and primer set.

| **Sample** | **Species** | **DAB1**<br>**PCR** | **DAB2**<br>**PCR** | **DAB3**<br>**PCR** | **DAB4**<br>**PCR** |
| --- | --- | --- | --- | --- | --- |
| B31737 | *N. elegans* | Y | Y | NA | NA |
| B31836 | *N. elegans* | Y | Y | NA | NA |
| B31991 | *Neopsephotus bourkii* | Y | Y | NA | NA |
| B32063 | *Neopsephotus bourkii* | Y | Y | NA | NA |
| B34717 | *N. pulchella* | Y | Y | NA | NA |
| B41912 | *N. pulchella* | Y | Y | NA | NA |
| B45495 | *N. chrysostoma* | Y | Y | NA | NA |
| B52124 | *N. splendida* | Y | Y | NA | NA |
| B52126 | *N. splendida* | Y | Y | NA | NA |
| Z11285 | *N. chrysostoma* | Y | Y | NA | NA |
| Z20835 | *N. chrysogaster* | Y | Y | Y | Y |
| Z3889 | *N. petrophila* | Y | Y | NA | NA |
| Z40386 | *N. chrysogaster* | Y | Y | Y | Y |
| Z40482 | *N. petrophila* | Y | Y | NA | NA |
| Z41397 | *N. chrysogaster* | NA | NA | Y | Y |
| Z3860 | *N. chrysogaster* | NA | NA | Y | Y |
| Z251 | *N. chrysogaster* | NA | NA | Y | Y |
| SB#<br>2035 | *N. chrysogaster* | NA | NA | Y | Y |
| 21-0168 | *L. Discolor* | Y | Y | Y | Y |

## Notes

### Competing Interest Statement

The authors have declared no competing interest.

